# Structural journey of an insecticidal pore-forming protein targeting western corn rootworm

**DOI:** 10.1101/2022.10.12.511876

**Authors:** G. Marini, B. Poland, C. Leininger, N. Lukoyanova, D. Spielbauer, J. Barry, D. Altier, A. Lum, E. Scolaro, C. Pérez Ortega, N. Yalpani, G. Sandahl, T. Mabry, J. Klever, T. Nowatzki, J-Z. Zhao, A. Sethi, A. Kassa, V. Crane, A. Lu, M.E. Nelson, N. Eswar, M. Topf, H.R. Saibil

## Abstract

Broad adoption of transgenic crops has revolutionized agriculture. However, resistance to insecticidal proteins by agricultural pests poses a continuous challenge to maintaining crop productivity and new proteins are urgently needed to replace existing transgenic traits. We identified an insecticidal membrane attack complex/perforin (MACPF) protein, Mpf2Ba1, with strong activity against western corn rootworm larvae and a novel site of action. By integrating X-ray crystallography, cryo-EM, and modelling, we determined monomeric, pre-pore and pore structures, revealing changes between structural states at atomic resolution. We discovered a monomer inhibition mechanism, a molecular “switch” associated with pre-pore activation/oligomerization upon gut fluid incubation and solved the highest resolution MACPF pore structure to-date. Our findings provide a mechanistic basis for Mpf2Ba1 effectiveness as an insecticidal protein with potential for biotechnology development.

**One-Sentence Summary:** The molecular mechanism of an insecticidal protein is revealed through 3D structures of the three main pore formation states

## Main Text

Pore-Forming Proteins (PFPs) are a major class of proteins that oligomerize into rings and undergo a large irreversible conformational change to punch holes in lipid bilayers, breaching the membrane of target cells. The unregulated ionic influx/efflux through the resulting pore causes a serious osmotic imbalance that eventually leads to cell necrosis (*1*, *2*).

PFPs derived from the entomopathogenic bacterium *Bacillus thuringiensis* (Bt) have been the basis for transgenic insect resistant crops, benefiting growers globally for almost three decades (*3*). *In planta* delivery of insect protection has proven to be highly effective and has environmental and economic benefits where deployed (*4*, *5*). One of the most devastating pests affecting maize production in North America and Europe and leading to significant yield loss is western corn rootworm (WCR) (*6*). Since the early 2000s, this pest has been effectively managed in North America by transgenic maize crops expressing Bt proteins from the Cry3 class (mCry3Aa, Cry3Bb1, eCry3.1Ab) or Gpp34Ab1/Tpp35Ab1 (previously known as Cry34Ab1/Cry35Ab1) (*7*, *8*). However, these proteins have encountered instances of reduced efficacy due to the development of resistance in field populations of WCR (*8*). Since the discovery rate of useful Bt-derived insecticidal proteins seems to be slowing, it is imperative to search for novel proteins from non-Bt organisms with high activity and new modes/sites of action (*8*, *9*).

We identified an insecticidal protein, Mpf2Ba1, active against WCR and carried out the first comprehensive structure-function characterization of a PFP from non-Bt organism, revealing key steps of pore formation. We demonstrate that gut fluid extracted from WCR larvae provides the necessary and sufficient environment for the protein activation, triggering conversion from the monomer to the pre-pore state and arming it for membrane penetration as it converts into the pore state. Using X-ray crystallography and cryo-EM, we reveal all three structural states.

Mpf2Ba1 is a 53.2 kDa protein isolated from *Pseudomonas monteilii* through a multi-step purification process informed by artificial diet bioactivity screening against WCR larvae. In artificial diet bioassays, purified recombinant Mpf2Ba1 and a variant bearing nine conservative mutations (var 1167; table S1) showed strong activity against WCR and northern corn rootworm (NCR) larvae, but no activity against other major agricultural pest species, suggesting that the protein has a certain degree of specificity to coleopteran insect pests (Fig. 1A and table S2). In addition, Mpf2Ba1 exhibited specific binding to WCR midgut tissue, caused rapid ion permeability in a southern corn rootworm cell line, suggesting pore formation, and provided strong root protection from corn rootworm damage when expressed in transgenic maize tested under greenhouse and field conditions (Fig. 1, B to D, fig. S1 and Supplemental text). Finally, artificial diet bioassays also show that Mpf2Ba1 is active against a field-derived population of WCR that exhibits signs of resistance to currently commercialized insecticidal proteins (mCry3A, and Gpp34Ab1/Tpp35Ab1; table S3 and Supplemental text). This indicates that Mpf2Ba1 elicits WCR mortality through a novel site of action and can bypass resistance against existing commercial trait proteins.

**Figure 1.**
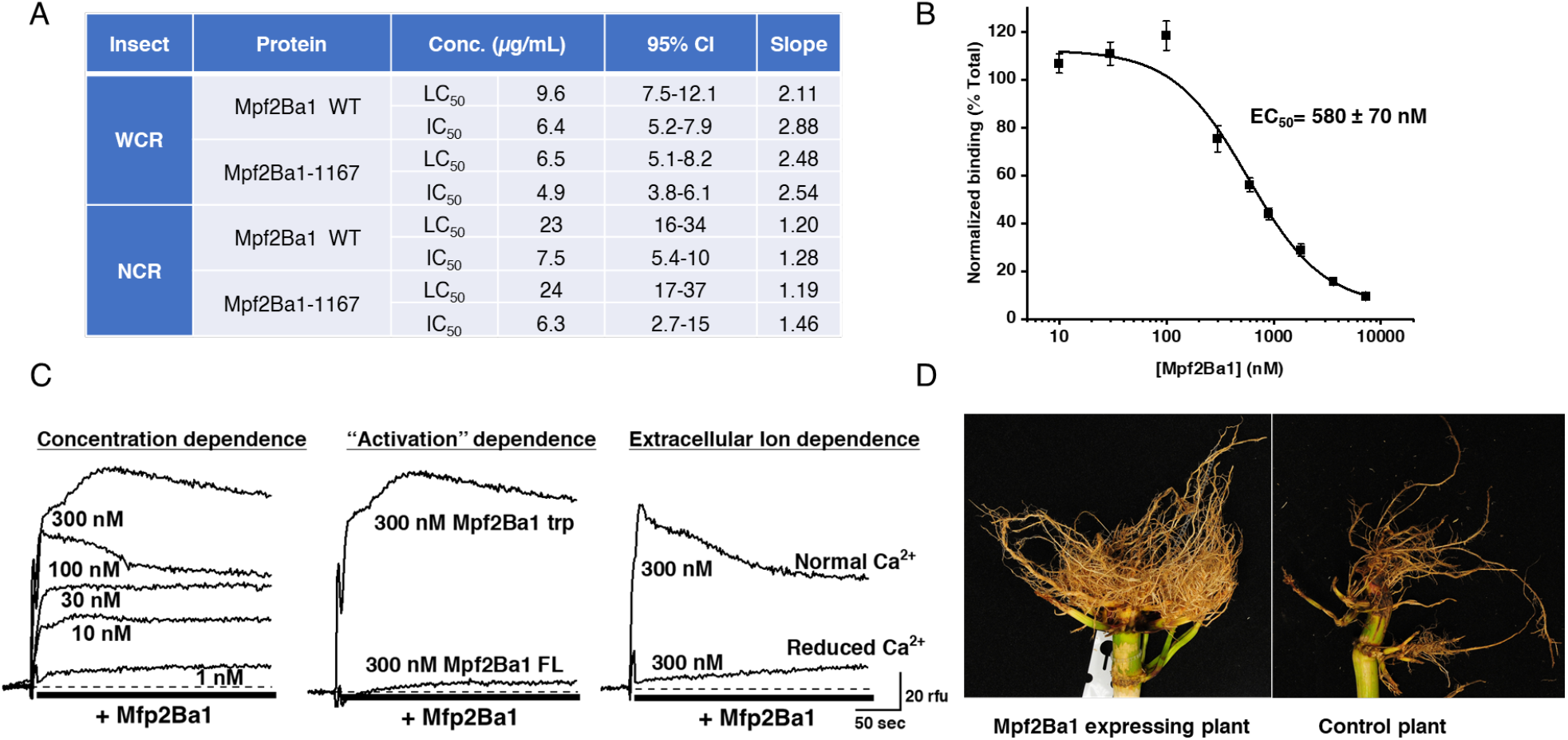
Activity of Mpf2Ba1 against corn rootworm. (**A**) Bioactivity of Mpf2Ba1 WT and Mpf2Ba1-1167 against WCR and NCR larvae in artificial diet bioassay. LC_50_ and IC_50_ (see Methods) are similar for both proteins with WCR exhibiting greater lethal sensitivity (lower LC_50_ value) than NCR. Confidence intervals (95% CI) and slope of the fitted curve from the analysis are also provided. (**B**) Specific binding of Mpf2Ba1 to WCR BBMVs is demonstrated by homologous competition of Alexa-Mpf2Ba1 by its unlabeled form. The values reflect the mean ± SEM from four observations of densitometry values determined from in-gel fluorescence images. (**C**) Real time fluorescence responses of southern corn rootworm cells (Du182Aa) loaded with the Ca^2+^-sensitive Fluo-3 dye are shown. The cells were exposed to Mpf2Ba1 WT at different concentrations after trypsin activation in normal media (left), to 300 nM Mpf2Ba1 before (FL) and after trypsin activation (trp) in normal media (middle), and to 300 nM Mpf2Ba1 after trypsin activation in normal Ca^2+^ or nominally Ca^2+^-free media (right). (**D**) Pictures of corn roots showing feeding damage by WCR from a field study that included plants expressing Mpf2Ba1and plants with the same genetic background that did not express Mpf2Ba1 (control plant).

We first solved the X-ray structure of Mpf2Ba1-1167 at 2.1 Å resolution (Fig. 2A). The protein is comprised of two domains – a N-terminal MACPF domain and a C-terminal β-prism domain. The MACPF domain has a central four-stranded antiparallel β-sheet with a characteristic ~90° bend. The β-sheet is interrupted by three insertions: two sets of transmembrane hairpins emerge from its base (TMH1, buried inside the hydrophobic core, and TMH2, facing the surface), and a helix-turn-helix (HTH) motif, harbouring several highly conserved residues in all known MACPF/CDCs (*10*) (fig. S2 and S3), intercalates at the bend of β4-strand. Both TMHs comprise a cluster of two α-helices and a loop, similar in length to the short TMHs of mammalian perforin-2 (PFN2, PDB IDs: 6U23 and 6SB3) (*11*, *12*).

**Figure 2.**
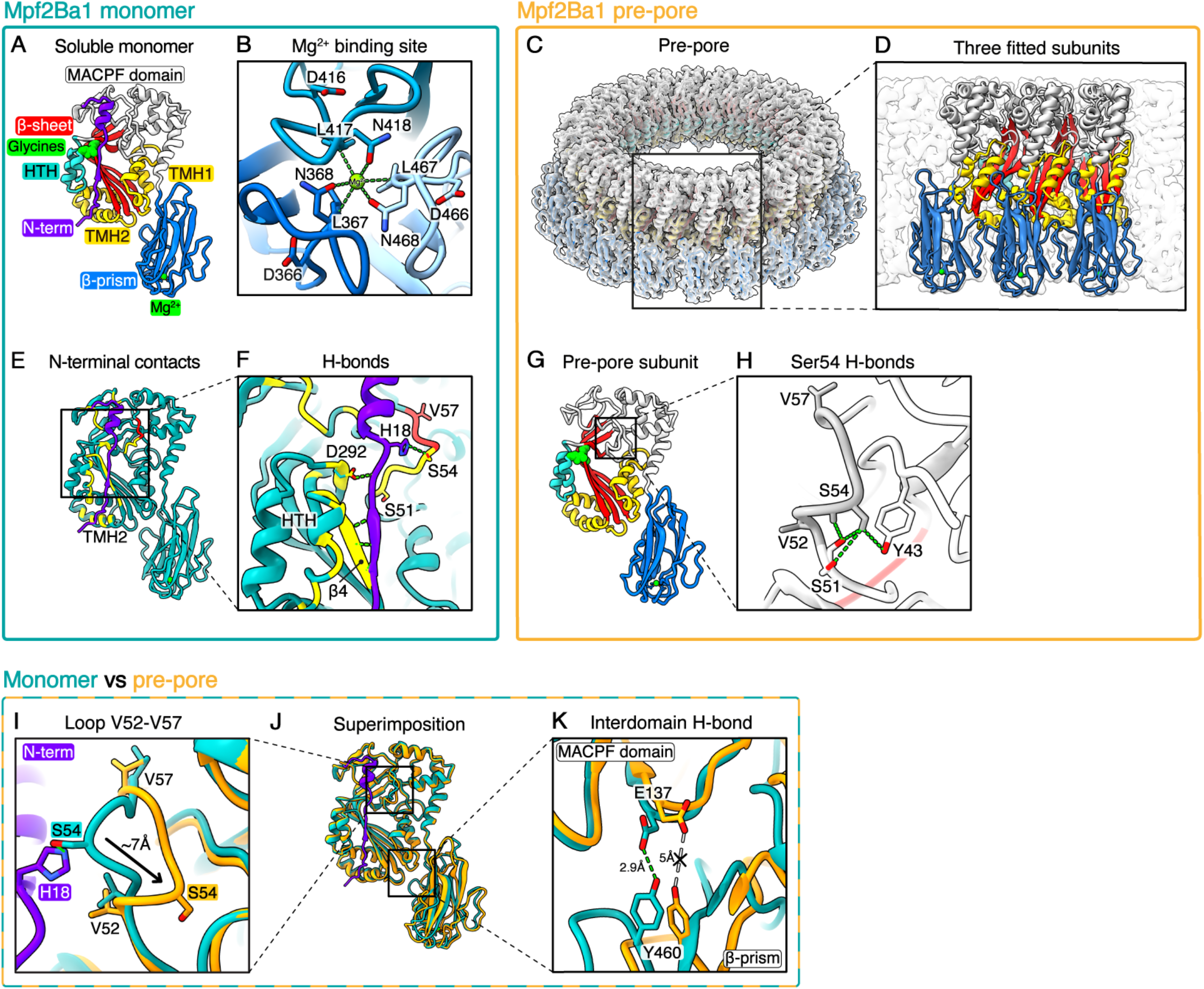
Structures of the Mpf2Ba1 monomer and pre-pore, and their comparison. (**A**) X-ray structure of Mpf2Ba1 soluble monomer. (**B**) View of the C-terminal membrane binding site showing conserved Asp, Leu and Asn (DLN) residues in the β-prism, and residues Leu and Asn coordinating a Mg^2+^ cation in the centre. (**C**) Single particle cryo-EM map of the 22-mer pre-pore at 3.1 Å resolution, with 22 fitted pre-pore subunits. **(D**) Inset of three pre-pore subunits fitted in the pre-pore map. (**E**) Interactions between the 30 N-terminal residues (in purple) and surrounding residues (in yellow) are highlighted. (**F**) Inset of the N-terminal contacts with surrounding residues (green dashes). (**G**) Model of the pre-pore subunit depicted according to the PFP domains. (**H**) Inset showing the buried position of loopVal52-Val57 in the pre-pore, where Ser54 contacts Tyr43 and Ser51, all conserved in bacterial MACPFs. (**I**) Inset showing comparison of loopVal52-Val57 positions between the soluble monomer (in cyan), and the pre-pore structure (in orange), where it is moved ~7 Å towards the MACPF domain core. (**J**) Superimposition of Mpf2Ba1 soluble monomer (in cyan) and pre-pore models (in orange). (**K**) Inset showing the ~15° rotation of the β-prism breaking the H-bond between the conserved Tyr460-Glu137.

The C-terminal domain adopts a β-prism fold, first identified as one of the three domains of insecticidal δ-endotoxins derived from Bt (*13*, *14*). It is structurally closely related to the C-terminal domain of Mpf1Aa1 from *Photorhabdus luminescens* (PDB ID: 2QP2, 26% seq. id; 1.58 Å RMSD over 114aa out of 147aa) (*15*) and, to a lesser extent, to the membrane associated β-prism domain of the multivesicular body subunit 12B (MABP-MVB12B) of human ESCRT-I complex (PDB ID: 3TOW, 21% seq. id; 1.6 Å RMSD over 84aa out of 147aa) (fig. S3) (*16*). In Mpf2Ba1, each of the three β-sheets of the β-prism, arranged around a local three-fold axis, contains a unique repeated motif: Asp-Leu-Asn (DLN). These DLN motifs create an octahedral geometry coordinating a Mg^2+^ at the symmetry axis through the Leu carbonyl oxygen and the Asn side-chain amide oxygen (Fig 2B and fig. S2). Similar domains of other MACPF proteins, such as Mpf1Aa1 (*15*), Mpf3Aa1, an insecticidal protein from *Chromobacterium piscinae* (PDB ID: 6FBM) (*17*), and PFN2 (*11, 12*), have Ca^2+^ as the coordinating ion. Structurally, the metal chelation seems to tuck the three β-prism loops into gaps between the adjacent β-sheets, increasing domain stability.

A major finding that facilitated Mpf2Ba1 structural characterization is that gut fluid extracted from WCR larvae is necessary and sufficient to induce oligomerization *in vitro* in the absence of brush-border membrane vesicles (BBMVs) (fig. S4). Upon incubation in gut fluid, an N-terminal cleavage after Lys25, as determined through Edman degradation sequencing, and subsequent oligomerization step are triggered by interaction of monomers with yet unidentified midgut component(s). Based on the observations that heat-inactivated gut fluid or the organic-extractable phase of gut fluid do not induce oligomerization (data not shown), we suspect that the oligomerization factor is a protein. Furthermore, we established that although N-terminal cleavage can be achieved using trypsin, WCR gut fluid is still necessary to trigger oligomerization. After gut fluid-induced oligomerization, Mpf2Ba1 monomers formed higher-order complexes that were isolated via size exclusion chromatography and imaged by cryo-EM. The 3D reconstruction of ring-shaped oligomers at 3.1 Å resolution revealed an intermediate state of Mpf2Ba1: a 22-mer pre-pore that is 80 Å in height, 260 Å in diameter and encircles a 120 Å-wide lumen (Fig. 2C and D, fig. S5 and table S4).

By fitting and refining the X-ray structure of Mpf2Ba1 into the pre-pore map, we discovered that 30 N-terminal residues are missing from the density, five residues more than the 25 identified biochemically (fig. S6). The missing density for those residues could be due to flexibility or disorder of the N-terminal region, or to additional proteolytic cleavage. Our finding that N-terminal cleavage is required for Mpf2Ba1 pore formation also on insect cells *in vitro* (Fig. 1C) prompted us to search for proteases active on the putative cleavage site before Thr31 observed in the pre-pore structure. Cathepsin-D and E emerged as the highest-ranked candidates, proteases that are abundant in midguts of WCR larvae (*18*), along with trypsin that would cleave at Lys25. Although further investigation is needed, cathepsins could play a role in triggering Mpf2Ba1 activation via N-terminal cleavage *in vivo*.

Comparison of the pre-pore structure with the soluble monomer shows small but significant conformational differences. In the monomer, the N-terminus lies adjacent to one side of the MACPF domain, forming a short β-strand–β-sheet interaction with the central β-sheet, and two more H-bonds: between Asp292, stabilized by the conserved Pro291, and the N-terminal backbone, and between one of the nitrogen atoms of N-terminal His18 imidazole ring and the oxygen atom of Ser54, highly conserved among bacterial PFPs (Fig. 2E and F, fig. S2). Upon N-terminal removal via gut fluid activation, these contacts are lost and residues on one side of the MACPF domain become fully exposed. Since we observed that some of these residues are involved in inter-molecular contacts with the neighbouring subunit, we think that the steric hindrance imposed by the N-terminus could help prevent premature interactions between soluble monomers.

The MACPF domain in the Mpf2Ba1 pre-pore remains mostly unchanged, except for loop Val52-Val57 that moves from an external position, in contact with the uncleaved N-terminus, to a buried position in the oligomeric structure, as measured from the accessible surface area per residue (Fig. 2G to I and fig. S7). The X-ray density for this loop is clear, albeit weaker and with higher B-factors than the surrounding residues, indicating greater flexibility. Because residues of this loop are in contact with the neighbouring subunit, it is likely that, upon N-terminal cleavage, this region rearranges and gets stabilized by the oligomeric conformation. Moreover, it seems that the loop is re-oriented ~7 Å to bring the highly conserved Ser54 from a solvent-exposed position in the monomer, in contact with His18 of the N-terminus, to a core-buried position towards the MACPF domain in the oligomer. There it engages in H-bonds with Tyr43 and Ser51 (Fig. 2H), also highly conserved in bacteria (fig. S2). The Ser54 “switch”, triggered by N-terminal cleavage and/or oligomerization, could possibly represent another key step in Mpf2Ba1 activation.

The pre-pore β-prism domain is rotated ~15° clockwise relative to its position in the monomeric structure. This slight rotation disrupts the only interdomain H-bond between MACPF and β-prism, involving the carboxyl oxygen of Glu137 and the hydroxyl group of Tyr460 (Fig. 2J and K, fig. S7 and Movie S1). Based on residue conservation, this H-bond seems specific to bacterial MACPFs (fig. S2), and its loss could be part of an allosteric signal relayed from the C-terminal domain upon membrane binding, possibly associated with oligomerization. Interestingly, although AlphaFold2 (*19*) accurately predicts Mpf2Ba1 domains and secondary structures of the soluble monomer and pre-pore, including the N-terminal α-helix, it does not predict either of these states correctly, as in both states the β-prism is not oriented precisely with respect to the MACPF domain, nor does it predict the pore state (fig. S6).

The transition from pre-pore to pore requires an energetically costly structural rearrangement (*20*). We were able to convert Mpf2Ba1 pre-pores into pores by either introducing lipids, in the form of POPC and 4% cholesterol mixed liposomes, or by raising the incubation temperature at ~53°C without adding lipids. When using liposomes, we observed membrane inserted structures that confirm the pore-forming activity of Mpf2Ba1 (Fig. 3A and fig. S4). Although heating is not the physiological trigger, it appears to catalyze the refolding of the α-helical TMHs into β-hairpins to form a β-barrel, which has a higher thermal stability than the helical conformation (*21*), confirming that pre-pore-to-pore conversion is endothermic (*22*). Heat-converted pores were assessed by negative-stain EM and optimized for cryo-EM single particle analysis.

**Figure 3.**
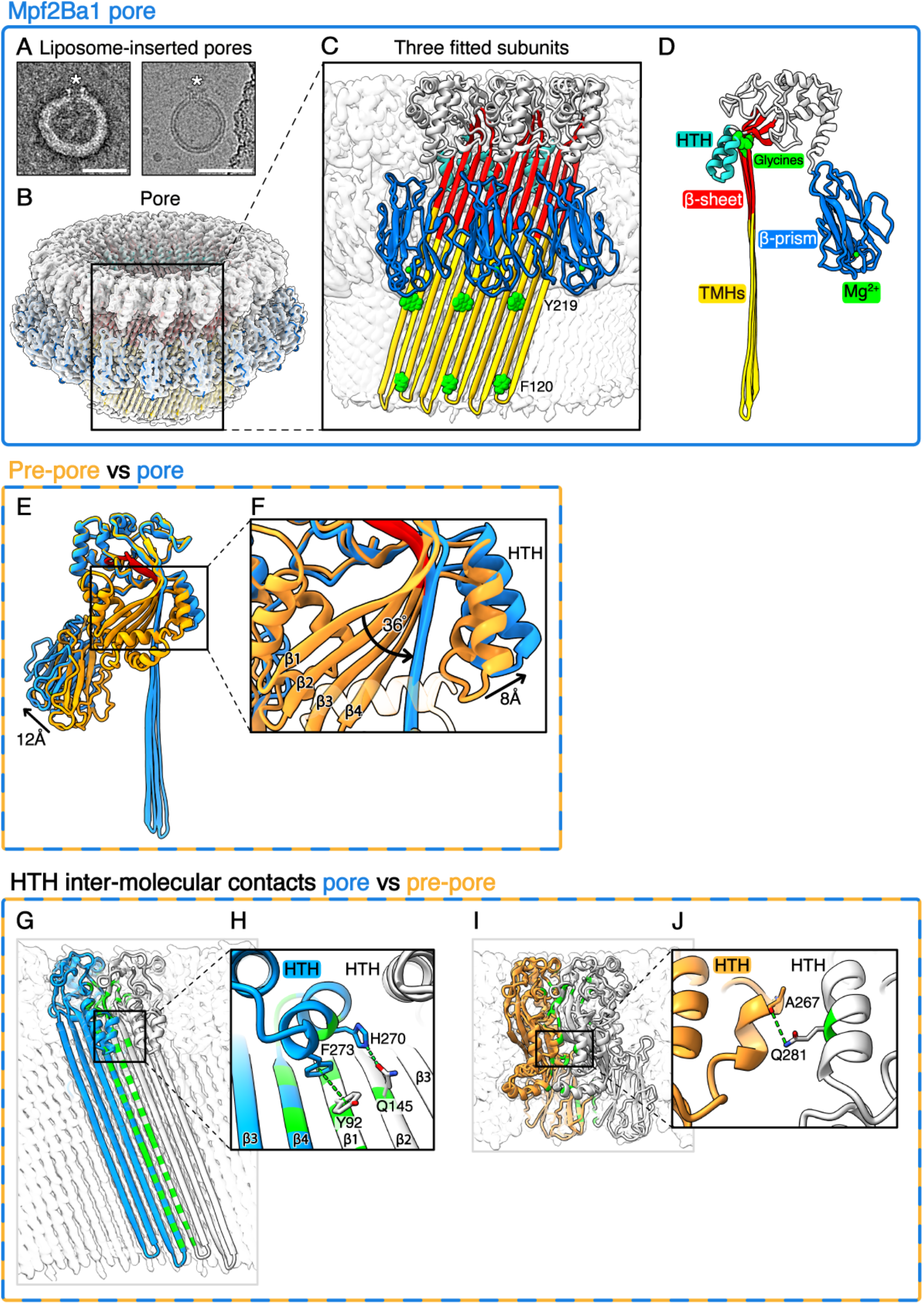
Mpf2Ba1 pore and comparison with the pre-pore structure. (**A**) Liposome-inserted pores (white stars) imaged in negative stain EM on the left, and cryo-EM on the right. Scale bars: 50 nm. (**B**) Single particle cryo-EM map of the 21-mer pore at 2.6 Å resolution and fitted pore subunits. (**C**) Inset of three pore subunits showing the aromatic residues Phe120 and Tyr219 (in green) forming an aromatic girdle around the β-barrel. (**D**) Model of the pore subunit. (**E**) Superposition of the pre-pore model (in orange) onto the pore model (in blue) revealing the conformational change between the two structures. The MACPF core signature motif is highlighted in red. (**F**) Inset showing the straightening of MACPF central sheet, where the strands β1-4 align towards the centre of the pore by an average dihedral angle of ~36°, and the HTH moves by 8 Å into the pore lumen. (**G**) Two neighboring pore subunits showing all intermolecular hydrogen bonds (in green). (**H**) View of the intermolecular contacts of HTH and strands β1 and β2 of the neighboring subunit. (**I**) Two neighboring pre-pore subunits showing all intermolecular H-bonds (in green). (**J**) View of the single intermolecular hydrogen bond between neighboring HTHs.

The 3D structure of the Mpf2Ba1 pore was determined at 2.6 Å resolution for the 21-mer, revealing a pore with outer diameter of 240 Å, β-barrel diameter of 135 Å, and a height of 120 Å, increased 40 Å from the pre-pore conformation (Fig. 3B, fig. S8 and table S4). The map resolution was appropriate to build an accurate atomic model by rigid body fitting of the Mpf2Ba1 pre-pore model without TMHs and subsequent manual building as β-hairpins, followed by full model refinement and validation (Fig. 3C and D, and table S5).

In the pore conformation, the β-sheet at the MACPF core straightens by an average dihedral angle of 36° to allow the helical bundles to refold into membrane-spanning β-hairpins and collectively form a β-barrel (Fig. 3E and F, and Movie S2). The flexibility for the central β-sheet straightening is provided by >95% conserved glycines among MACPF/CDCs, that function as a hinge (fig. S2 and S3) (*15*, *24*).

The straightening of the MACPF central β-sheet repositions the HTH, which is pushed ~8 Å towards the lumen of the pore, restricting the diameter locally from 135 Å to 110 Å (Fig. 3F). This shift brings the HTH in close lateral contact with residues of the neighbouring subunit, where Phe273 contacts Tyr92 on β1-strand in a π-π interaction, and the side-chain nitrogen of His270 forms a H-bond with the oxygen of Gln145 on β2-strand (Fig. 3G and H). A single H-bond was detected in the pre-pore, connecting neighbouring HTHs (Fig. 3I and J). In pleurotolysin MACPF domain (PlyB), the HTH displacement was proposed as one of the key triggers for the pore conformational change (*23*). In Mpf2Ba1 we see that HTH motion follows the straightening of the TMHs and does not move relative to them (Movie S3), but the HTH intermolecular contacts change throughout pore formation. Based on the different interactions, we propose key regulatory roles for HTH in: (i) preventing premature interaction between monomers in solution, (ii) controlling the transition from pre-pore to pore, and (iii) stabilizing inter-subunit interactions in the final pore structure (*23*–*25*).

The biggest conformational change during pore formation involves the two TMHs that unfurl from α-helical bundles into ~100 Å-long membrane-spanning β-hairpins, a unique feature of the MACPF/CDC superfamily (Fig. 3B–E, Movies S2 and S3) (*26*). As in other MACPF structures, each subunit of Mpf2Ba1 pore contributes four β-strands to the β-barrel (84 β-strands for Mpf2Ba1 21-mer). The β-strands are tilted by 20° from the membrane normal, in agreement with the β-barrel shear number *S* = *n*/2 found in giant β-barrel architecture (*27*, *28*). The β-hairpins are stabilized by intra- and intermolecular H-bonds between β-strands, and the β-barrel is stabilized inside the membrane by aromatic residues, frequently found in transmembrane β-barrels oriented towards the polar phospholipid moiety and commonly reported as aromatic girdles (*29*–*31*). In Mpf2Ba1, the polar residues Phe120 on β2-strand at the intracellular side of the membrane bilayer, and Tyr219 on the β3-strand at the extracellular side, are probably responsible for positioning and stabilizing the β-barrel inside the membrane. Lastly, the C-terminal β-prism domain moves ~12 Å outwards relative to its pre-pore position (Fig. 3E and fig. S7), possibly to allow the MACPF domain to approach and fully span the membrane.

In addition to inter-subunit interactions, the non-uniform surface charge distribution of PFP oligomers seems to contribute to subunit stabilization and membrane insertion. Indeed, neighbouring Mpf2Ba1 pre-pore and pore subunits showed complementary charge interaction, as inferred from electrostatic potential maps. On one side of the MACPF domain, a negatively charged bulge and a positively charged pocket match exactly with a positively charged pocket and a negatively charged bulge on the neighbouring subunit in both pre-pore and pore. Moreover, the charge distribution on the inner and outer surfaces of the pore is as expected for a transmembrane protein, with a highly polar lumen-facing surface, mostly water exposed, and a hydrophobic membrane-facing surface of the β-barrel, forming an aliphatic nonpolar belt of approximately ~40 Å in height (fig. S9).

In conclusion, we have identified and characterized the non-Bt insecticidal protein Mpf2Ba1, providing mechanistic details for how it effectively kills the devastating maize pest WCR by forming pores in midgut membranes targeting a different binding site than the currently deployed insecticidal proteins (Fig. 4). We solved the Mpf2Ba1 X-ray structure and showed how N-terminal residues inhibit the association of monomers in the soluble form. We then demonstrated that proteolysis in the presence of gut fluid activates monomers that self-assemble into pre-pores, which we reconstructed by cryo-EM at 3.1 Å resolution. The subtle but significant changes between monomers and pre-pore subunits revealed molecular details related to target recognition and activation/oligomerization. The heat-induced conversion of pre-pores into pores allowed us to solve the 21-mer pore cryo-EM structure at 2.6 Å resolution, showing the dramatic conformational change of the pore subunits an their HTH-mediated inter-subunit stabilization.

**Figure 4.**
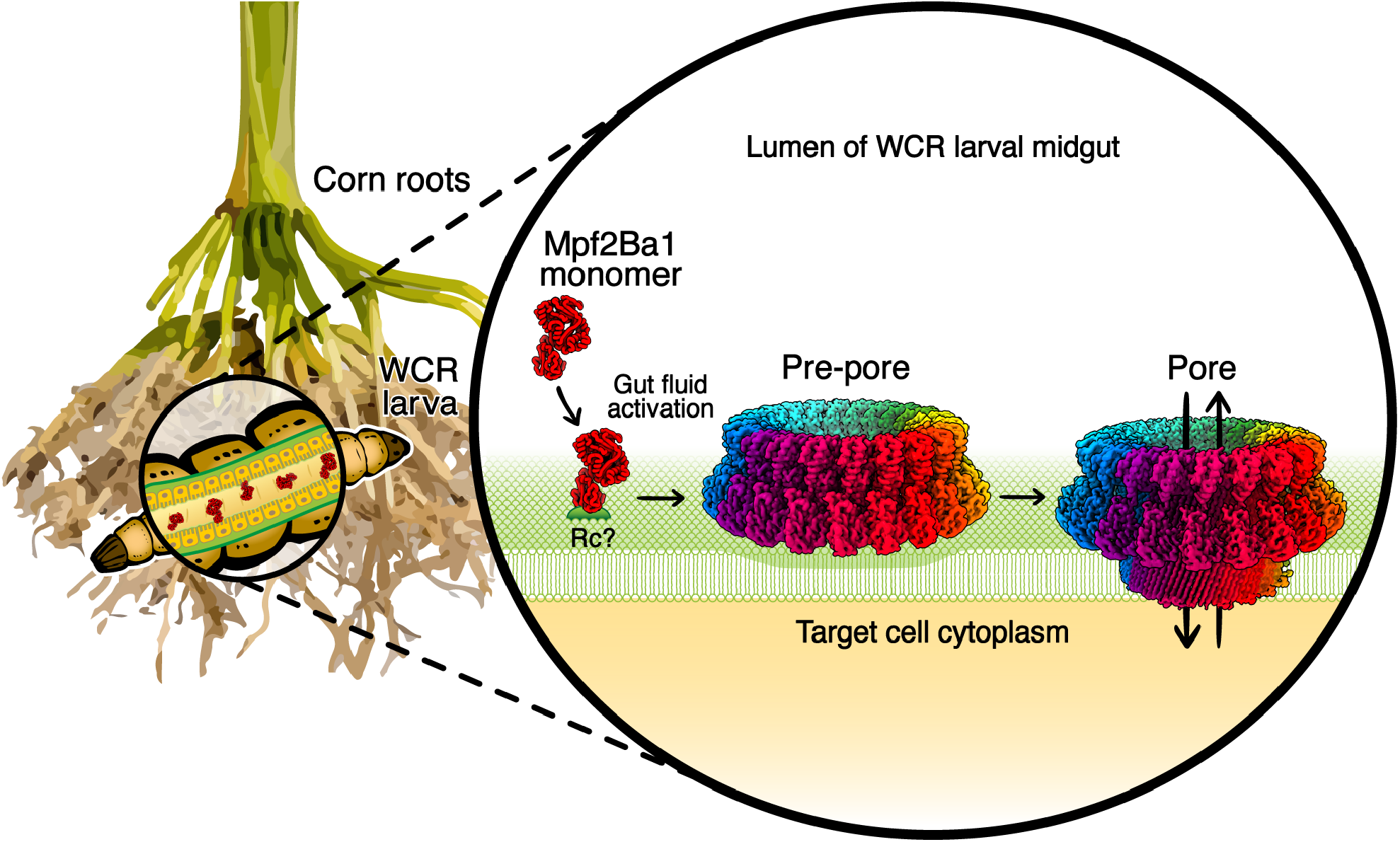
Model of Mpf2Ba1 in-plant protection and insecticidal activity against WCR larvae. The model shows the three Mpf2Ba1 conformations characterized in our study in the context of the WCR larval midgut. The soluble monomers expressed by plant root tissues reach the larval midgut and are activated by gut fluid before engaging a not-yet-identified receptor (Rc) on the membrane of the target cells where they oligomerize into pre-pores. Pre-pores convert into irreversible transmembrane pores that allow uncontrolled ion flux across the plasma membrane, disrupting cellular function and causing the larvae to die of starvation. For the preservation of maize yield, management of WCR pest by proteins with novel sites of action is urgently needed, and Mpf2Ba1 represents an excellent candidate for this purpose.

Our work provides not only the first high-resolution oligomeric structures for a MACPF insecticidal protein targeting WCR, but also the highest-resolution MACPF pore structure to date. We demonstrated that Mpf2Ba1 efficacy and unique site of action give it great potential for protecting maize from WCR damage, particularly where there is resistance to current trait proteins.

## Acknowledgments

We thank the Protein Core Facility, Crop Transformation Systems and Controlled Environments Group of Corteva Agriscience for help with protein expression, plant transformation, and greenhouse support, respectively; USDA-ARS, Plant Sciences Institute, Beltsville, MD for providing IPLB-DU182A cells; Z. Hou for help in structural analysis; D. Cerf for bioassay of *D. speciosa*; C. Stewart, X-ray facility manager and T. P. Stewart, electron and light microscopy facility manager at Iowa State University; D. Houldershaw, S. Malhotra, T. Cragnolini, A. Sweeney and L. Genz for IT and modeling support, C. Bagneris for support in the Rosalind Franklin Lab; S. Chen for imaging support and all members of Topf, Saibil and Orlova groups for fruitful scientific discussion at Birkbeck College, University of London.

## Author contributions

M.N., N.E., M.T. and H.R.S. conceived the idea. G.M., B. P., C.L., A. Lum, T.M., J.K., J-Z.Z., A.K., E.S., and C.P-O. performed the experiments, C.L., D.S., J.B., D.A., N.Y., M.N. and N.E. provided the samples, B.P. performed the X-ray crystallography, G.M. and N.L. performed cryo-EM, G.M. performed the single particle analysis and molecular modelling, A.K. conducted the diet bioassays, J-Z.Z. and A.S. planned and conducted the experiments with field derived resistant WCR colonies, V.C. supervised vector construction, G.S. and T.M. conducted the greenhouse experiments, J.K. and T.N. planned and conducted the field experiments, A.Lu provided supervision and guidance, G.M., B.P., M.N., N.E., M.T., and H.R.S. wrote the manuscript.

Conceptualization: M.N., N.E., M.T., and H.R.S.

Methodology: G.M., B.P., C.L., M.N., N.E., M.T., and H.R.S.

Investigation: G.M., B.P., C.L., N.L., D.S., J.B., D.A., A.Lum, C.P-O, N.Y., G.S., T.M., J.K., T.N., J-Z.Z., A.S., A.K., E.S., M.N., N.E., M.T., and H.R.S.

Visualization: G.M., B.P., C.L., N.L., M.N., N.E., M.T., and H.R.S.

Funding acquisition: A.Lu, M.N., N.E., and H.R.S.

Project administration: G.M., M.N., N.E., M.T., H.R.S.

Supervision: V.C., A.Lu, M.N., N.E., M.T. and H.R.S.

Writing – original draft: G.M., B.P., M.N., N.E., M.T., and H.R.S.

Writing – review & editing: G.M., M.N., N.E., M.T., and H.R.S.

## Data and materials availability

sequences for Mpf2Ba1 and Mpf2Ba1-1167 were deposited in NCBI under accession numbers, OP537915 and OP575916, respectively. Cryo-EM maps of Mpf2Ba1 pre-pore and pore are deposited with the Electron Microscopy Database under accession numbers EMD-15882 and EMD-15883. PDB coordinates of Mpf2Ba1 are deposited with the Protein Data Bank under accession numbers 8B6U, 8B6V and 8B6W for monomer, pre-pore and pore subunits.

## Supplementary Materials

Materials and Methods

Supplementary Text

Figs. S1 to S9

Tables S1 to S5

References (*31*–*83*)

Movies S1 to S3

