## Supplementary material for "Structural journey of an insecticidal pore-forming protein targeting western corn rootworm"

**This PDF file includes:**

Materials and Methods
Supplementary Text
Figs. S1 to S9
Tables S1 to S5
Thumbnails and captions for Movies S1 to S3

**Other Supplementary Materials for this manuscript include the following:**

Movies S1 to S3

#### Materials and Methods

Identification, isolation, and purification of Mpf2Ba1

The insecticidal protein Mpf2Ba1 was identified through activity-based screening and protein purification as described in Schellenberger et al., 2016 and Wei et al., 2018 (*32*, *33*). Briefly, insecticidal activity against western corn rootworm (WCR) was measured with a cell lysate of bacterial strain JH34071-1 that was grown in Tryptic Soy broth (TSB, peptone from casein 15 g/L; peptone from soymeal 5.0 g/L; NaCl 5.0 g/L) and cultured 1 day at 28° C with shaking at 200 rpm. Bioassays with WCR were conducted by mixing the bacterial cell lysate with *Diabrotica* diet (Frontier Agricultural Sciences, Newark, Del.) in a 96 well format. WCR neonates were placed into each well of a 96 well plate. The assay was run for four days at 25° C and then was scored for insect mortality and stunting of insect growth. The scores were noted as dead "3", severely stunted "2" (little or no growth but alive), stunted "1" (growth to second instar but not equivalent to controls) or normal growth "0". Samples demonstrating mortality or severe stunting were studied further. Once the activity was established, the equivalent sample was subjected to heat and Pronase treatments and resubmitted for insect bioassays to confirm that the insecticidal activity was due to a proteinaceous component.

To identify the bacterial strain where activity was observed, genomic DNA of isolated strain JH34071-1 was prepared according to a library construction protocol developed by Illumina and sequenced using the Illumina® Genome Analyzer IIx (Illumina Inc., 9885 Towne Center Drive, San Diego, Calif. 92121). The nucleic acid contig sequences were assembled and open reading frames were generated. The 16S ribosomal DNA sequence of strain JH34071-1 was BLAST™ searched against the NCBI database (*34*), identifying strain JH34071-1 as a *Pseudomonas monteilii* strain. To isolate and identify the active protein, cell pellets of strain JH34071-1 were homogenized at -30,000 psi after re-suspension in 20 mM Tris buffer, pH 8 with cOmplete™, EDTA-free protease inhibitor cocktail (Roche, Indianapolis, Ind.). The crude lysate was cleared by centrifugation and brought to 75% saturation with ammonium sulfate. The 75% ammonium sulfate solution was then centrifuged, and the supernatant was discarded. The pellet portion was suspended in 20 mM Tris pH 8.0 and then brought to 1.5 M ammonium sulfate with the addition of a 2 M ammonium sulfate, 20 mM Tris pH 8.0 solution. This solution was clarified and loaded onto a TSKgel™ Phenyl- 5PW column (Tosoh Bioscience, Tokyo, Japan) equilibrated in 20 mM Tris pH 8.0, 1.5 M ammonium sulfate. Insecticidal activity eluted with a gradient to 20 mM Tris, pH 8. Active fractions were pooled, concentrated on 10 kDa molecular weight cutoff centrifugal concentrators (Sartorius Stedim, Goettingen, Germany) and desalted into 20 mM piperazine pH 9.5 using a Sephadex G25 (GE Healthcare, Piscataway, N.J.) column. The desalted pool was loaded onto a Mono Q™ column (GE Healthcare, Piscataway, N.J.) equilibrated in 20 mM piperazine, pH 9.5 and eluted with a gradient of 0 to 0.4 M NaCl. Active fractions were pooled and loaded onto a Superdex™ 200 column (GE Healthcare) equilibrated in phosphate buffered saline (PBS). SDS-PAGE analysis of fractions indicated that WCR activity coincided with a prominent band after staining with GelCode™ Blue Stain Reagent (Thermo Fisher Scientific). The protein band was excised, digested with trypsin and analyzed by nano-liquid chromatography/electrospray tandem mass spectrometry (nano-LC/ESLMS/MS) on a Thermo Q Exactive™ Orbitrap™ mass spectrometer (Thermo Fisher Scientific, 81 Wyman Street, Waltham, Mass. 02454) interfaced with an Eksigent™ NanoLC™ 1-D Plus nano-Ic system (AB Sciex™, 500 Old Connecticut Path, Framingham, Mass. 01701). Protein identification was done by internal database searches using Mascot® (Matrix Science, 10 Perrins Lane, London NW3 1QY UK). The candidate protein was designated originally as IPD090Aa (Mpf2Ba1). Cloning and recombinant expression confirmed the insecticidal activity of the Mpf2Ba1 polypeptide (accession # OP537915) against WCR. Note that for preparing plant expression constructs (see below) the *mpf2ba1* gene sequence was modified to include an additional Ala immediately following the translation initiation Met, thus increasing the amino acid reference numbers by one. Except as noted, all of the structure and biochemistry studies described below used this protein sequence.

Generation of Mpf2Ba1-1167

Simulated gastric fluid (SGF) is a common method for assessing the potential allergenicity of nonfood proteins (*35*). It was observed that Mpf2Ba1 was stable when exposed to SGF (data not shown). An engineered SGF-susceptible variant, Mpf2Ba1-1167 (accession # OP575916), was generated by making five phenylalanine to tyrosine and four isoleucine to leucine mutations (table S1). Mpf2Ba1-1167 had increased susceptibility to SGF exposure while maintaining wild type potency against WCR (Fig. 1). Mpf2Ba1-1167 was used as a representative of Mpf2Ba1 for crystal structure determination.

Mpf2Ba1 and Mpf2Ba1-1167 $LC_{50}/IC_{50}$ determinations

Mpf2Ba1 or Mpf2Ba1-1167 protein was incorporated into artificial diet to conduct bioassays on key rootworm species (WCR, NCR, and *Diabrotica speciosa*). Corn rootworm diet was prepared according to manufacturer's guidelines for *Diabrotica* diet (Frontier, Newark, DE). The test involved six different protein doses plus buffer control with 32 observations for each dose in each bioassay. Neonate larvae were infested into 96-well plates containing a mixture of the Mpf2Ba1 proteins (5 μL/well) and diet (25 μL/well), each well with approximately 5 to 8 larvae (<24 h post hatch). After one day a single larva was transferred into each well of a second 96-well plate containing a mixture of the Mpf2Ba1 (20 μL/well) and diet (100 μL/well) at the same concentration as the treatment to which the insect was exposed on the first day. The plates were incubated at 27°C, 65% RH in the dark for 6 days. The 50% lethal concentration ($LC_{50}$) for polypeptides in the bioassay was calculated using "Dose Response Add-In for Excel" based on Probit analysis. Larvae were scored on a scale of 0-3 (unaffected, stunted, and severely stunted, respectively) or dead. Dead and severely stunted counts were pooled as total response for the calculation of the concentration for inhibition of 50% of the individuals ($IC_{50}$) using the same method (Fig. 1A).

Greenhouse efficacy of Mpf2Ba1-expressing plants

Greenhouse efficacy results for events generated from Mpf2Ba1 constructs are shown in figure S1. Note that plant expression constructs utilized a modified *mpf2ba1* gene sequence that included a codon for an additional Ala immediately following the translation initiation Met. Events were generated using 13 different constructs designed to produce a range of Mpf2Ba1 accumulation. Constructs were introduced into *Agrobacterium tumefaciens* strain LBA4404, and those transformants were used to infect immature embryos of the maize inbred line PHR03 as described (*36*). Efficacy for events derived from all 13 constructs was observed relative to negative control events (Empty) as measured by root protection from western corn rootworm. Root protection was measured according to the number of nodes of roots injured (CRWNIS = corn rootworm node injury score) using the method developed by Oleson, et al. (2005) (*37*). The root injury score is measured from "0" to "3" with "0" indicating no visible root injury, "1" indicating 1 node of root damage, "2" indicating 2 nodes or root damage, and "3" indicating a maximum score of 3 nodes of root damage. Intermediate scores (e.g. 1.5) indicate additional fractions of nodes of damage (e.g. one and a half nodes injured).

Field efficacy of Mpf2Ba1-expressing plants

Field efficacy results for Mpf2Ba1 expressing plants are shown in figure S1B. Events analyzed in this study were generated such that Mpf2Ba1 accumulated preferentially in the plastids of roots. In both Construct 1 and 2, Mpf2Ba1 transcript was driven by the SB-RCC3-PCOA118532A promoter (SEQ ID #12) (*38*) followed by the ZM-HPLV9 INTRON1 (SEQ ID #8) (*39*). Transcript generation in both was terminated by the OS-T30 terminator (SEQ ID #31) (*40*). The two constructs differed only in the transit peptide enabling plastid localization. In Construct 1, Mpf2Ba1 was targeted to plastids by fusing the Mpf2Ba1 transcript (minus the first Met codon) at the 5' end to nucleotides encoding: `MCGSTPRTAKFIRAPHLGVKAWKMVAWPGSARWAKRLSMGPVPVYSAWNQAPMTASMARRPFLISLVR`. In construct 2, the transit peptide was: `MLIISSTPHLGVKAWKMVAWPGSARWAKRLSMGPVPVYSAWNQAPMTASMARRPFLISLVR`.

The experimental unit in the field was a single-row plot of corn 3 m in length and a row spacing of 76 cm. The experimental design was a randomized complete block with subsamples, planted at 8 locations, with treatments randomized within each of 3 replications. Treatments included 3 Mpf2Ba1 transformation events from Construct 2 (9 plots per location), 2 Mpf2Ba1 transformation events from Construct 1 (6 plots per location), 2 entries of the commercial event DAS-59122-7 as the positive control (6 plots per location), and 2 entries with no events for corn rootworm (CRW) protection as the negative control (6 plots per location). Treatments were evaluated from 4 locations that sustained sufficient root injury to the negative control (at least 0.75 nodes of injury). Additional experimental constructs not related to Mpf2Ba1 were included in the experiment but are not reported. The commercial event DAS-59122-7 expresses the Gpp34Ab1/Tpp35Ab1 proteins from *Bacillus thuringiensis* strain PS149B1 that act together as a binary insecticidal protein to provide protection against corn rootworm CRW larvae (*41*). All treatments were tested in a single hybrid with the same genetic background. A seed treatment containing the insecticide, thiamethoxam, at a rate of 0.25 mg a.i./kernel (Cruiser® 250; Syngenta Crop Protection, Inc., Greensboro, NC, USA) was applied to seeds in all treatments. This is the labeled rate for control of certain secondary insect pests of corn but does not control CRW.

The source of infested WCR eggs was a non-diapausing colony maintained by the Corteva Insectary Production Research group located in Johnston, IA. Root injury to the treatments was evaluated after the peak period of CRW larval feeding had occurred at each location. Roots were evaluated by digging a sub-sample of 3 roots per plot, washing the root systems clean of soil, and then visually assessing the amount of CRW larval injury (node-injury score) using the Iowa State 0-3 node-injury scale (*37*).

A linear mixed model was applied to model node-injury scores across locations. Data for node-injury score ($Y_{ijmnks}$) of location $(L)_i$, replication $(R)_j$, construct $(P)_m$, event $(E)_n$, plot $(K)_k$ and plant $s$, were modeled as a function of an overall mean $\mu$, factors for location, location by replication, construct, event, location by construct, location by event, plot within each location $(K/L)_{ik}$ and a residual within each location $(\varepsilon/L)_{ijmnks}$. The model can be specified as:

$$Y_{ijmnks} = \mu + \underline{L_i} + \underline{(L \times R)_{ij}} + P_m + \underline{E_n} + \underline{(L \times P)_{im}} + \underline{(L \times E)_{in}} + \underline{(K/L)_{ik}} + \underline{(\varepsilon/L)_{ijmnks}}$$

where construct was treated as fixed effect, and all the other effects were treated as independent normally distributed random variables with means of zero. *F*-tests were used to assess significance for fixed effects. *T*-tests using standard errors from the model were conducted to compare treatment effects. A difference was considered statistically significant if the *P*-value of the difference was less than 0.05. All data analysis and comparisons were made in ASReml 3.0 (VSN International, Hemel Hempstead, UK, 2009).

Mpf2Ba1 mode of action

To understand the mechanism of Mpf2Ba1 toxicity, specific binding of the purified protein with WCR midgut tissue was evaluated by *in vitro* competition assays. Midguts were isolated from third instar WCR larvae to prepare brush border membrane vesicles (BBMVs) as described in Zhao et al., 2016 (*42*) using amino-peptidase activity to track enrichment. BBMVs represent the apical membrane component of the epithelial cell lining of insect midgut tissue and therefore serve as a model system for how insecticidal proteins interact within the gut following ingestion.

Purified Mpf2Ba1 was diluted to 1 mg/ml and processed with immobilized trypsin resin (Sigma, T1763 or Promega V901B) using resin at 1:1 (v:v) ratio incubated at room temperature overnight. An aliquot of the trypsin-processed protein was then labeled with Alexa-Fluor® 488 (Life Technologies) and unincorporated fluorophore was separated from labeled protein using buffer exchange resin (Life Technologies, A30006) following manufacturer's recommendations. Prior to binding experiments, proteins were quantified by gel densitometry following Simply Blue® (Thermo Fischer Scientific) staining of SDS-PAGE resolved samples that included BSA as a standard.

To establish specific binding and to evaluate binding affinity, BBMVs (2.5 μg) were incubated with Alexa-labeled Mpf2Ba1 (Alexa-Mpf2Ba1; 5 nM) in binding buffer (50 mM sodium chloride, 2.7 mM potassium chloride, 8.1 mM disodium hydrogen phosphate, and 1.47 mM potassium dihydrogen phosphate, pH 7.5 containing 0.1% Tween20®; 100 μL) for 1 hr at RT in the absence and presence of increasing concentrations of unlabeled Mpf2Ba1 (0.01-7.2 μM). Centrifugation at 20,000×g was used to pellet the BBMVs to separate unbound Alexa-Mpf2Ba1 remaining in solution. The BBMV pellet was then washed twice with binding buffer to eliminate remaining unbound Alexa-Mpf2Ba1. The final BBMV pellet (with bound fluorescent protein) was solubilized in reducing Laemmli sample buffer, heated to 100 °C for 5 minutes, and subjected to SDS-PAGE using 4-12% Bis-Tris polyacrylamide gels (Life Technologies). The amount of Alexa-Mpf2Ba1 in the gel from each sample was measured by a digital fluorescence imaging system (ImageQuant™ LAS4000 - GE Healthcare). Digitized images were analyzed by densitometry software (Phoretix™ 1D, TotalLab, Ltd.). Figure 1B shows that Mpf2Ba1 binds specifically to WCR BBMVs. The densitometry values were normalized to the signal observed for Alexa-Mpf2Ba1 binding in the absence of competitor ("Total") and plotted vs the concentration of unlabeled protein present during competitions. The resulting distribution was then fit to a logistic equation in OriginPro software (ver. 2021B; Originlab Corp.); y = A2 + (A1-A2)/(1 + (x/x0)^p), where A1 and A2 are the maxima and minima, respectively, x is the unlabeled protein concentration, and x0 and p correspond to the $EC_{50}$ and slope values, respectively.

Pore formation assay

To evaluate the ability of Mpf2Ba1 to undergo pore formation, a cell line derived from *Diabrotica undecimpunctata* (southern corn rootworm) designated IPLB-Du182A (Du182A) was used (*43*). Du182A cells were maintained in culture by growing in SF-900 II medium (Gibco,

10902) supplemented with 3% heat inactivated FBS (Gibco, 16140) and antibiotics (Gibco, 5240) in T-75 flasks at 27° C. To detect pore formation, Du182A cells were grown in 96-well assay plates (Corning, 3594). After reaching approximately 70% confluence, the medium was removed, and the cells were washed with serum-free medium. The cells were then loaded with Fluo-4 AM (10 μM from DMSO stock; Thermo Fisher Scientific, F14201) in medium consisting of 50% Hank's buffered saline (HBSS; Gibco 14025092) supplemented with 3.7 mM $CaCl_2$ (final concentration of 5 mM) and 50% serum-free SF-900 II medium along with 2.5 mM probenecid (Sigma, P8761) and 0.2% (v:v) Pluronic F-127 (Thermo Fisher Scientific, P6867) for 1 hr at 27° C. The loading solution was then removed and replaced with assay medium consisting of 50% HBSS supplemented with $CaCl_2$ (5 mM final concentration), 50% serum-free SF-900 II and 2.5 mM probenecid. The loaded cells were then placed in a Flexstation 3 (Molecular Devices) along with a compound plate that contained appropriate working stock dilutions of Mpf2Ba1 proteins. Mpf2Ba1 (1.6 mg/ml) was activated by treating the full-length protein with immobilized trypsin beads (Sigma T1763) 1:1 (v:v) at 25°C until no full-length protein was detected (>30 hours). Testing for pore formation was achieved by transfer of activated Mpf2Ba1 protein (unless otherwise stated) from the compound plate to the assay plate using the Flexstation's built-in fluidics module and real time monitoring of fluorescence intensity controlled by SoftmaxPro (v5.2, Molecular Devices) running in Flexmode. The transfer of Mpf2Ba1 was carried out after monitoring baseline fluorescence for 30 seconds (sampling interval ~1 sec) and the cell responses were monitored for an additional 4.5 minutes. To construct figures, data were exported into OriginPro (v.2021b; OriginLabs) where baseline fluorescence was subtracted from each well and the subtracted responses were plotted versus elapsed time.

Edman degradation sequencing

N-terminal cleavage was determined through Edman degradation sequencing, as described previously (*44*). Untreated Mpf2Ba1 (2 μg) and Mpf2Ba1 processed with WCR gut fluid incubated at 37° C overnight (3 μl of 3 mg/ml Mpf2Ba1 mixture) were separated on a 4-12% Bis-Tris Gel and then transferred to a PVDF membrane using an IBlot® (ThermoFisher) semi-dry transfer system. The membrane was then stained with Sypro™ Ruby (ThermoFisher) to visualize protein bands and bands corresponding to untreated Mpf2Ba1, monomeric Mpf2Ba1, and oligomeric Mpf2Ba1 following gut fluid treatment were cut from the membrane and subjected to sequencing. Both the monomer and oligomeric bands revealed cleavage of 25 amino acids before Ala26 in Mpf2Ba1.

Cross-resistance to commercial trait proteins

The activity of Mpf2Ba1 and Mpf2Ba1-1167 were evaluated in artificial diet bioassays using a WCR population collected from a research location in Readlyn, IA, which is situated within an area where there is high use of commercial hybrids expressing proteins that control damage from rootworms (Cry3 class and Gpp34Ab1/Tpp35Ab1). This population of WCR has been designated the "Readlyn" strain and has been shown to have signs of resistance to both mCry3A and Gpp34Ab1/Tpp35Ab1 (*45*). With this in mind, the insecticidal activities of the Mpf2Ba1 proteins against the Readlyn population were compared to their activity against a susceptible non-diapausing laboratory population of WCR. The format of the bioassays was similar to the method described above for bioactivity assessments against susceptible WCR larvae with certain differences. There were five to eight protein concentrations plus buffer control with 24 observations for each dose. Mortality data were analyzed following PROC PROBIT procedure in SAS software (SAS Institute, 2013) for obtaining $LC_{50}$ and 95% confidence interval.

Crystal growth and data collection

Crystals of Mpf2Ba1-1167 were grown by hanging drop vapor diffusion method at 22°C. Crystals of Mpf2Ba1-1167 were obtained by a 1:1 ratio of 10 mg/ml protein solution and reservoir solution containing 0.2M $MgCl_2$ hexahydrate, 0.1M HEPES pH=7.5 and 30% PEG 400. Crystals were flash frozen in liquid $N_2$ and mounted on a Rigaku Micromax-007 HF X-ray source at Iowa State University Macromolecular X-ray Crystallography facility. Data were collected using an R-Axis IV++ image plate detector. Mpf2Ba1-1167 crystals diffracted to 2.1 Å and belong to space group $P4_12_12$ with one molecule in the asymmetric unit. Diffraction data were indexed and integrated with iMOSFILM (CCP4) and scaled with SCALA (CCP4).

Structure determination of Mpf2Ba1-1167

The atomic structure of Mpf2Ba1-1167 was solved using the molecular replacement program PhaserMR (CCP4). The structure of the Mpf1Aa1protein from *Photorhabdus luminescens* (PDB ID: 2QP2) (*15*) was used as the search model. A suitable solution for the rotation and translation functions was identified. The sequence for the Mpf2Ba1-1167 was then built into the electron density using WinCoot© (*46*). The model was refined using Refmac5 from CCP4 (*47*) to an R-factor=0.236 and R-free=0.267 with >96% of amino acids in allowed regions of the Ramachandran Plot.

Sequence analysis

The degree of Mpf2Ba1 sequence conservation in bacteria was obtained using the ConSurf server (*48*). ConSurf defines a conservation score based on the evolutionary rate of a particular position through alignment and phylogenetic analysis of homologous sequences. A total of 249 sequences from a BLAST™ (*34*) search in the nonredundant database were used to derive normalized conservation scores. The conservation scores were distributed into nine grades, from the most variable 1 to the most conserved 9. The generated alignment file was read into UCSF ChimeraX (*49*) and represented on the Mpf2Ba1 structure, with the corresponding residues colored according to the scores (fig. S2).

The sequence/structure conservation for Mpf2Ba1 was obtained running HHPred (*50*), a method for sequence database searching and structure prediction very sensitive in finding remote homologs, on the PDBmmCif30 database, based on the Protein Data Bank (PDB) (*51*) and with a maximum sequence identity of 30%, for the sequence of the MACPF domain (residues 31-331) and for the β-prism domain (residues 338-484) separately.

Oligomer formation and purification for structure determinations

Soluble monomeric Mpf2Ba1, purified at 6 mg/mL stock in 1x PBS buffer (Gibco, 10010056), was diluted with an equal volume of freshly thawed WCR gut fluid (GF). GF was prepared from freshly dissected larval guts (third instar) mixed with 1x PBS (5 μL/gut) incubated on ice for 90 mins and then centrifuged at 20k x g for 15 at 4° C to collect the supernatant. The Mpf2Ba1/GF mixture was incubated for 1-4 hours at 37˚C with rotary mixing. To purify oligomers after WCR GF treatment, the incubated suspension was loaded onto a Superose 6 Increase 10/300 GL size exclusion column (GE Healthcare Biosciences, Piscataway, NJ, USA), pre-equilibrated with 1x PBS pH 7.4 (Gibco, 100100) filtered using a 0.22 μm filter and degassed. Protein was eluted with the same buffer at 0.5 mL/min and fraction size 0.5 mL for 1.5 column volumes. For vitrification and cryo-EM imaging, fractions containing Mpf2Ba1 pre-pores were pooled and concentrated to ~0.8 mg/mL using a centrifugal filter 100 kDa cutoff (SigmaAldrich).

Mpf2Ba1 pores were obtained by incubating purified pre-pores, or a 1:1 mixture of Mpf2Ba1 soluble monomers and GF, with liposomes (described below) for 1 hour. Alternatively, pores were obtained by sonication in a water bath or by incubation at ~50˚C for 1 hour without adding lipids, the latter yielding the best pre-pore to pore conversion. Subsequent size exclusion chromatography onto a Superose 6 Increase 10/300 GL size exclusion column (GE Healthcare Biosciences, Piscataway, NJ, USA), elution buffer as described above separated pores from monomers and/or bigger liposomes, if present. Fractions containing Mpf2Ba1 pores were pooled and concentrated to ~0.6 mg/mL using a centrifugal filter 100 kDa cutoff (SigmaAldrich) and then vitrified and imaged by cryo-EM.

Liposome preparation

Liposomes were prepared from a mixture of 1-palmitoyl-2-oleoyl-glycero-3-phosphocholine (POPC 16:0-18:1; 10mg/mL Avanti Polar Lipids, Inc., Alabaster, Alabama) and 4% plant derived cholesterol (100 mg; Avanti Polar Lipids) in chloroform. The lipid mixture at 1 mg/mL was dried under a stream of nitrogen gas in a glass vial (Avanti Polar Lipids). The resulting thin film was rehydrated in 1x PBS buffer pH 7.4 (Gibco). The lipid suspension was vortexed, sonicated in an ultrasonic water bath for 10 minutes at ~40˚C (Branson 1510 Ultrasonic Cleaner) and flash frozen in liquid nitrogen and thawed at least 3 times. The lipid mixture was then extruded through polycarbonate nanopore filters of 100 nm and 80 nm pore size (Avanti Polar Lipids) to generate large unilamellar liposomes of ~100 nm, using an Avanti Mini-Extruder with on a 40˚C heated plate (Grant Bio).

Negative stain EM

Carbon coated grids (EMS, 200 copper mesh) were cleaned and hydrophilized by glow discharge for 30 s (Pelco easiGlow, TedPella) at 25 mA before application of 4 µL of Mpf2Ba1 sample, diluted to 0.1 mg/mL. After 1 min waiting time, excess liquid was blotted away with filter paper (Whatman #1) and the sample was stained with 3 µL droplets of 2% uranyl formate (Genereon #24762-5). Images were acquired on a Tecnai F20 electron microscope (Thermo Fisher Scientific) with a DE20 camera (Direct Electron) at ×30,000 magnification (2.05 Å/pixel) and 0.75–1.5 µm underfocus.

Cryo-EM sample preparation

Freshly purified Mpf2Ba1 pre-pores concentrated to ~0.8 mg/mL and pores concentrated to ~0.6 mg/mL (3 µl) were applied to graphene oxide coated holey carbon grids (EMS, CFlats R 1.2/1.3 and 2/1 300 copper mesh) prepared following the procedure of Cheng et al., 2020 (*52*). After 5 s blotting time, samples were plunge-frozen in liquid ethane cooled by liquid nitrogen (*53*), using a Vitrobot Mark IV (Thermo Fisher Scientific) and stored under liquid nitrogen until use. Screening of cryo-EM conditions was performed on a 200 kV F20 (Thermo Fisher Scientific) with a DE-20 direct electron detector camera (DirectElectron).

Data collection and image processing of pre-pores

A dataset of 3578 movies was collected on 300 kV Titan Krios microscope (Thermo Fisher Scientific) with a K2 Summit direct electron detector camera and Quantum energy filter (Gatan).

Image processing and single particle analysis were performed in cryoSPARC v2.9.0 (Structura Biotechnology) (*54*), unless otherwise stated (fig. S3). Electron micrograph movie frames were aligned using cryoSPARC patch motion correction. CTF parameters were estimated, refined, and corrected using cryoSPARC patch CTF estimation. Particles were picked and subjected to several rounds of two-dimensional (2D) classification, selecting classes with identifiable secondary structure features. After manual curation to exclude low-quality micrographs based on poor CTF resolution (>9 Å) and particles with deviated local motion trajectories, a total of 64,428 particles were selected and used to generate an *ab initio* three-dimensional (3D) model without imposed symmetry (C1). This map was used as initial model for homogeneous refinement. D22 symmetry was imposed in the refinement of the pre-pore, since particles, 2D class averages and *ab initio* 3D model showed 22-mer pre-pores paired in dimers. The final refinement was carried out using non-uniform refinement with per-particle CTF refinement. The final map was sharpened using DeepEMhancer 'tightTarget' option from the unfiltered unmasked halfmaps (*55*). The resolution of the map was determined with masking-effect corrected Fourier Shell Correlation (FSC) as implemented in cryoSPARC. Local resolution was estimated in the final map using 'blocres' (*56*) (implemented in cryoSPARC). The pre-pore image processing workflow is summarized in table S4.

<u>Data collection and image processing of pores</u>

A dataset of 21,728 movies of pore structures was collected on a 300 kV Titan Krios electron microscope with a K3 direct electron detector and Quantum energy filter (Gatan) and processed in cryoSPARC v3.2.0 (Structura Biotechnology) (*54*), unless otherwise stated (fig. S7). Electron micrograph movie frames were aligned by MotionCor2 (*57*) in 5x5 patches. CTF parameters were estimated, refined, and corrected using CTFFIND4 (*58*). A round of manual curation was used to exclude sub-optimal micrographs based on poor CTF resolution (>8 Å). Particles were picked and subjected to several iterations of two-dimensional classification in cryoSPARC v3.2.0 (Structura Biotechnology) (*54*), selecting only the most homogeneous class averages showing identifiable secondary structure features. 2D classes showed a distribution of oligomer stoichiometries, ranging from the largest 24-fold (7% of end view particles), 23-fold (11%), 22-fold symmetry (52%) to 21-fold (30%), demonstrating heterogeneity of oligomerization. It was evident from raw side-views and 2D classes that Mpf2Ba1 pores also dimerized tail-to-tail. 1,413,104 selected particles contributed to an *ab initio* 3D model without imposed symmetry (C1), which was used as initial model for homogeneous refinement and 3D classification. The dimers of tail-to-tail pores had the barrel of one pore inserted inside the other and two different stoichiometries between inner and outer pore (fig. S6, 3D classes red and blue). Despite the most common stoichiometry in the 2D classes being a 22-mer, in the 3D reconstructions the inner ring of each dimer showed a clear and consistent 21-fold symmetry, while the outer ring was a mixture of 22-fold and higher symmetries. To best refine the inner 21-fold symmetric pore while excluding density from the poorly resolved outer ring in the dimer, a mask enclosing only the 21-mer pore of the dimer was created from the full 3D map, using the segmentation tool SEGGER (*59*) in UCSF Chimera (*60*). Local refinement, coupled to per-particle CTF refinement and with C21 symmetry imposed, was run to iteratively refine the masked sub-volume of the Mpf2Ba1 21-mer. The local resolution of the final 3D reconstruction was calculated using 'blocres' in cryoSPARC with FSC threshold 0.5. The resolution of the map was determined using masking-effect corrected Fourier Shell Correlation (FSC) in cryoSPARC v3.2.0 (Structura Biotechnology) (*54*). The 3D refined final map was sharpened using deepEMhancer 'highRes' option from the unfiltered unmasked halfmaps (*55*). The pore image processing workflow is summarized in table S4.

Model building, analysis, and validation

Pre-pore model: the X-ray structure of Mpf2Ba1 was fitted into the cryo-EM map after initial rigid body assignment using RIBFIND (*61*) and successive flexible fitting using Flex-EM (*62*), both from CCP-EM suite (*63*). Model refinement was performed in Coot v0.8.9.2 (*46*) and ISOLDE (*64*).

Pore model: the pre-pore model was rigid body fitted into the cryo-EM pore map using the same programs above. Transmembrane β-hairpins were manually built into the β-barrel densities using Coot v0.8.9.2 (*46*) and iteratively refined using ISOLDE (*64*) and, to ensure proper geometry, real space refinement in PHENIX v1.19.2 (*65, 66*).

For the N-terminal cleavage site protease identification, Procleave_sequence webserver (*67*) was used providing the sequence of the cleavage site.

The electrostatic potential of a single subunit of the Mpf2Ba1 pore was calculated using PDB2PQR software (*68*), to prepare the model PDB file for continuum solvation in a .pqr file that was used as input for the DelPhi suite (*69*) to solve the Poisson-Boltzmann equation for the pore subunit.

For both pre-pore and pore models, Q-score from MapQ (*70*), TEMPy SMOCf (*62*, *71*), MolProbity (*72*) and Strudel (*73*) scores were used to validate the final model (table S5).

Model-to-map FSC was estimated using PHENIX v1.19.2 (*66*). Structure visualization, fitting of multiple subunits, segmentation, analysis of charges and contacts, images and movies were generated with both UCSF Chimera (*60*) and ChimeraX (*49*).

#### Supplementary Text

Oligomerization of Mpf2Ba1

As stated in the main text, proteolytic activation of Mpf2Ba1 is necessary, but not sufficient to induce oligomerization. Oligomerization was first observed with incubation of the protein with gut fluid extracted from WCR larvae. However, it was not clear whether the N-terminal truncation *per se* was triggering oligomerization or not. Therefore, Mpf2Ba1 was treated with trypsin and chymotrypsin to see if these proteases truncated the protein and whether oligomerization was observed. Both proteases truncated Mpf2Ba1 to a fragment that was very similar in size to the one observed with gut fluid processing, but neither caused oligomer formation to occur (not shown). Trypsin was then used to activate Mpf2Ba1 and the resulting truncated peptide at Ala26 was exposed to gut fluid where oligomerization was observed. Furthermore, gut fluid treated with protease inhibitors could drive oligomer formation of truncated Mpf2Ba1 indicating that additional proteolysis was not needed for oligomerization to occur. These observations indicated that an additional factor must be present in gut fluid that interacts with activated Mpf2Ba1 and triggers oligomerization. Finally, incubation of trypsin-truncated Mpf2Ba1 with BBMVs (not shown) or with southern corn rootworm cells (Du182Aa) also resulted in oligomer/pore formation (Fig. 1B and C), indicating that binding to a membrane receptor triggers oligomerization.

Activity of Mpf2Ba1 against resistant strains of WCR

To demonstrate that Mpf2Ba1 works through a mechanism that is distinct compared to the WCR traits that are currently commercialized, we compared its activity against a laboratory susceptible population of WCR to its activity against a field-collected population from Readlyn, IA, an area where there is high use of commercialized traits (denoted as 'Readlyn'). The Readlyn population has been shown to have signs of resistance to both mCry3Aa and Gpp34Ab1/Tpp35Ab1 (*45*).

Using an artificial diet, we performed concentration-response bioassays and computed the $LC_{50}$ values for Mpf2Ba1 against the susceptible rootworm population and the "Readlyn" population (Table S3). The Readlyn population is much less sensitive to both Gpp34Ab1/Tpp35Ab1 and mCry3A with resistance ratios ($LC_{50}$ against Readlyn population/$LC_{50}$ against susceptible population) of 8- and 43-fold, respectively. Mpf2Ba1 or Mpf2Ba1-1167 shows a resistance ratio of <1-fold, indicating that there is no cross-resistance to mCry3Aa or Gpp34Ab1/Tpp35Ab1. These results show that the mechanism by which Mpf2Ba1 elicits mortality is different compared to either mCry3Aa or Gpp34Ab1/Tpp35Ab1.

Field efficacy

For evaluation of Mpf2Ba1-expressing plants under field conditions, second-generation (T2) non segregating hybrid maize seeds, derived from two constructs transformed to contain Mpf2Ba1 (two events from Construct 1 and three events from Construct 2) were evaluated at 4 locations across the U.S. corn belt in 2016 (Johnston, IA; Mankato, MN; Brookings, SD, and Mansfield, IL).

The field trials were conducted on land that contained late planted conventional corn in the previous season to attract corn rootworm beetles for egg laying and to enhance natural infestations. In addition, plots at all locations were manually infested with 750 to 1,500 WCR eggs per plant (depending on location) between growth stages V2 and V4. Based on mean ± SD node-injury scores obtained from the negative control plants, corn rootworm larval feeding pressure was classified as high at Johnston, Mankato, and Brookings (2.9±0.23, 2.7±0.33 and 1.8±0.68, respectively); and moderate at Mansfield (1.5±0.45).

General observations from all locations indicated that the predominant corn rootworm species was the WCR. Combining the data across all four testing locations, plants expressing the Mpf2Ba1 protein provided very good root protection from feeding injury by corn rootworm larvae (fig. S1B). Details on statistics for the fixed and random effects are summarized in the Material and Methods section. Mean node-injury scores between the 2 Mpf2Ba1 constructs were numerically lower, but not significantly different from the root protection provided by the commercial hybrid corn line DAS-59122-7 across the four locations ($P < 0.05$; fig. S1B). Both Mpf2Ba1 constructs provided good protection compared with the negative control ($P < 0.05$, Table 1). As reference, a node-injury score of 1.0 under field conditions has been estimated to cause a 15 to 18% reduction in corn grain yield (*74, 75*).

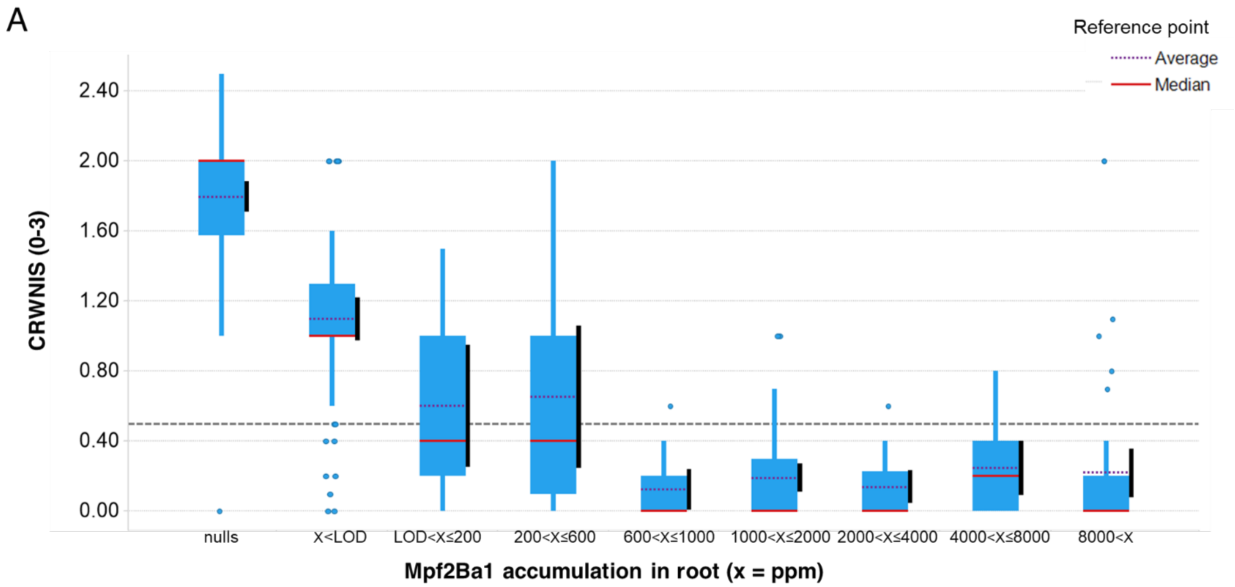


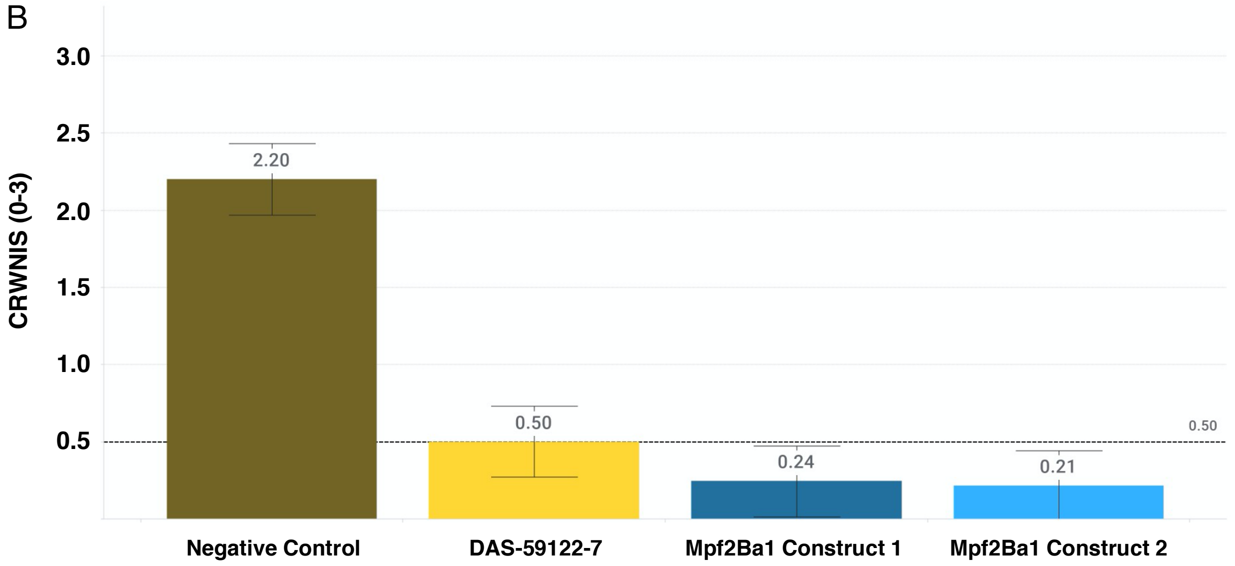


**Fig. S1. Mpf2Ba1 root expression vs. corn root worm nodal injury score (CRWNIS).** (**A**) *In planta* efficacy data expressed as the corn rootworm node injury score (*37*) as a function of protein expression (mass spectrometry) in the root determined using the peptide sequence EPTPPGYTK as described previously in Hu and Owens, 2011 (*76*). Plants expressing Mpf2Ba1 at 600-1000 ppm show preferred levels of rootworm control (CRWNIS < 0.5). LOD is limit of detection by mass spectrometry. (**B**) Field efficacy data comparing plants expressing Mpf2Ba1 (two different constructs) to DAS-59122, the commercial event that expresses Gpp34Ab1/Tpp35Ab1, and untreated control hybrid plants (Negative Control). The bars reflect data collected from four different locations in 2016.

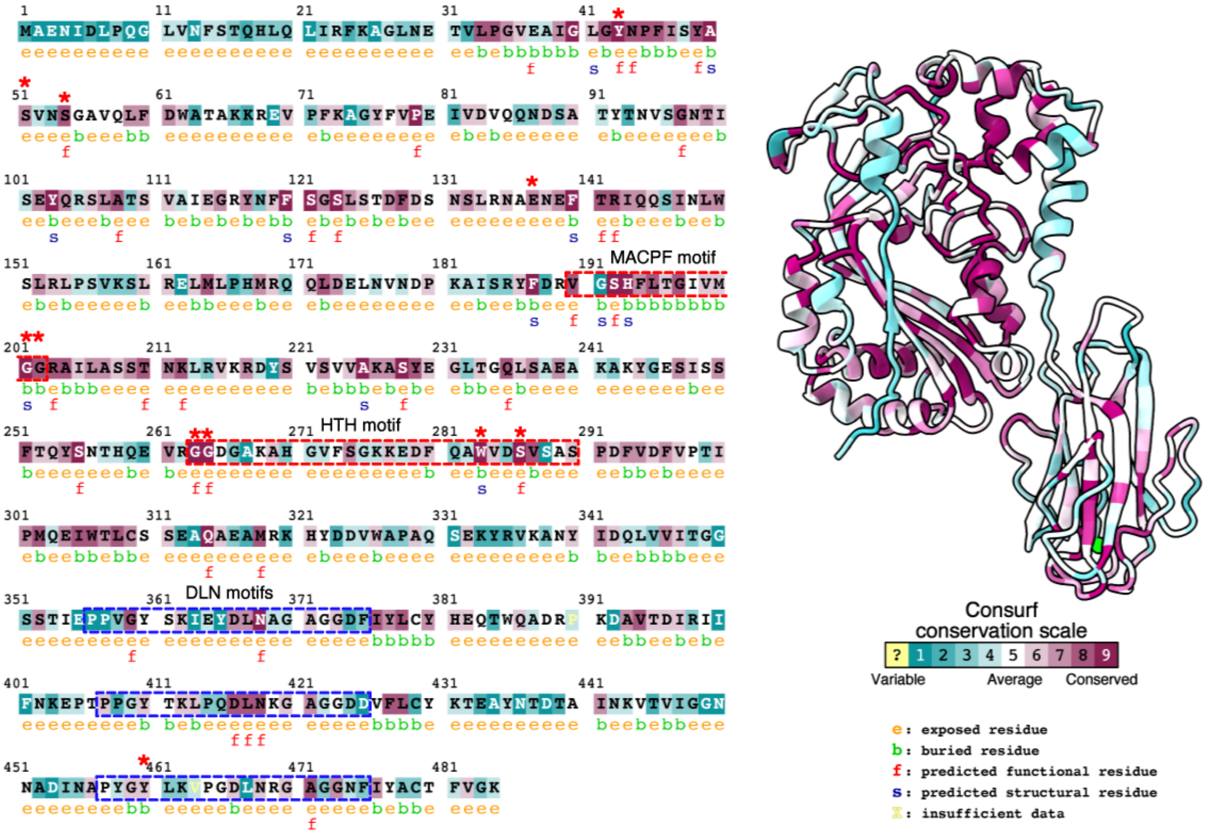


**Fig. S2. Mpf2Ba1 sequence conservation and residue evolution among bacteria.** Each sequence position of Mpf2Ba1 is colored according to conservation scores obtained using ConSurf (*48*), to show residue conservation and evolutionary rate. Red stars highlight key conserved residues: glycines 201-202, in the MACPF central sheet, glycines 263-264 and Trp283 (*10*) in the HTH motif, and the conserved H-bond between Glu137 and Tyr460. Red dashed boxes frame the sequences of MACPF signature motif Y/W-G-T/S-H-F/Y-$X_6$-GG (*15, 77*) which in Mpf2Ba1 diverges slightly from the consensus with a non-conserved change from Tyr/Trp to Val ($^{190}$V-G-S-H-$X_{6(FLTGIVM)}$-GG$^{202}$), and the sequence of the HTH motif; blue dashed boxes frame the sequences of Mpf2Ba1 C-terminal repeated MABP motif PxG(Y/F)(T/S)xIxxDLNxxxxGx(Y/F); see also Fig. 3E (in red). The X-ray crystal structure has been rendered by color coding attribute using ChimeraX (*49*) to reflect the evolutionary conservation.

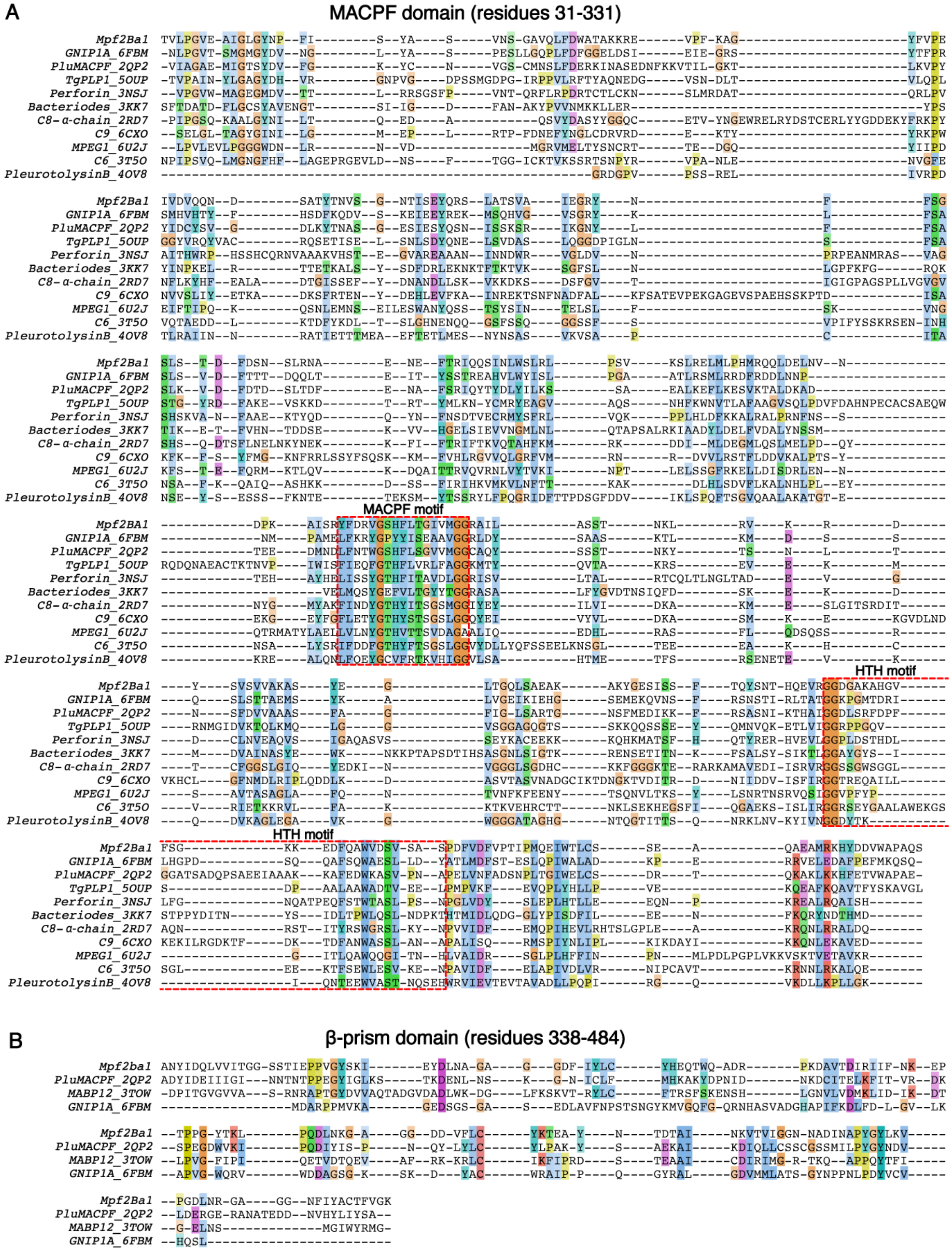


**Fig. S3. Sequence alignment and residue conservation between MACPF/CDCs.** (**A**) Mpf2Ba1 MACPF domain sequence aligned to the closest structural homologues using HHPred (*50*) and visualized in Jalview (*78*): Mpf3Aa1 (GNIP1Aa, PDB ID: 6FBM) (*17*), Mpf1Aa1 (Plu-MACPF from *Photorhabdus luminescens*, PDB ID: 2QP2) (*15*), *Toxoplasma gondii* perforin-like protein 1 (PDB ID: 5OUP) (*79*), human perforin (PDB ID: 3NSJ) (*80*), MACPF *Bacteroides thetaiotaomicron* (PDB ID: 3KK7) (*10*), human C8 protein (PDB ID: 2RD7) (*81*), mouse complement component-9 (PDB ID: 6CXO) (*82*), MPEG1/Perforin-2 (PDB IDs: 6U23 and 6SB3) (*11*, *12*), human C6 protein (PDB ID: 3T5O) (*83*), and pleurotolysin B (PDB ID: 4OV8) (*23*). Red stars highlight key conserved residues: glycines 201-202, in the MACPF central sheet, glycines 263-264 and Trp283 (*10*) in the HTH motif, and the conserved H-bond between Glu137 and Tyr460. Red dashed boxes frame the sequence of MACPF signature motif Y/W-G-T/S-H-F/Y-$X_6$-GG (*15*, *77*), which in Mpf2Ba1 has a non-conserved change from Tyr/Trp to Val ($^{190}$V-G-S-H-$X_{6(FLTGIVM)}$-GG$^{202}$), and the sequence of the HTH motif. (**B**) Mpf2Ba1 β-prism domain sequence aligned to the closest structural homologues MACPF/CDCs: Mpf1Aa1 (PDB ID: 2QP2) (*15*), Mpf3Aa1 (PDB ID: 6FBM) (*17*), and the multivesicular body subunit 12 MABP domain of Human ESCRT-I complex (PDB ID: 3TOW) (*16*).

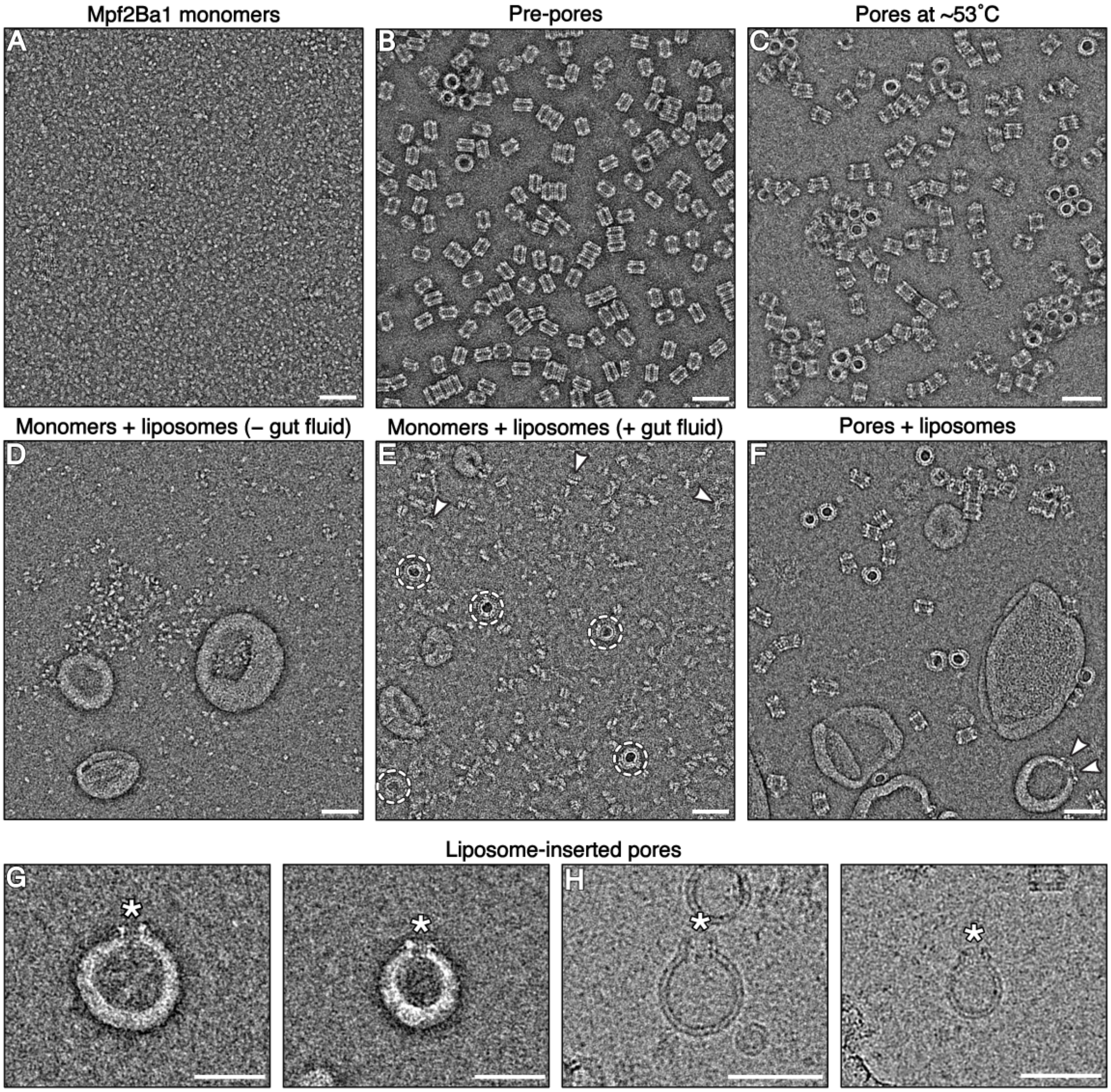


**Fig. S4. Negative stain EM to assess pore sample preparation.** The micrographs show Mpf2Ba1 (**A**) soluble monomers alone; (**B**) pre-pores oligomerized from monomers incubated in WCR gut fluid; (**C**) pores converted from pre-pores using the heating method without liposomes/lipids; (**D**) monomers incubated in POPC and 4% cholesterol liposomes without gut fluid, showing no oligomerization; (**E**) monomers activated by gut fluid and incubated in liposomes, showing weak oligomerization into pores (white dashed circles) and small arc-like assemblies, (white arrowheads); (**F**) pores converted from pre-pores after incubation in liposomes, (white arrowheads indicate liposome-inserted pores). (**G, H**) Magnified views of liposome-inserted Mpf2Ba1 pores (white stars) imaged by negative stain (**G**) and cryo-EM (**H**). Scale bars: 50 nm.

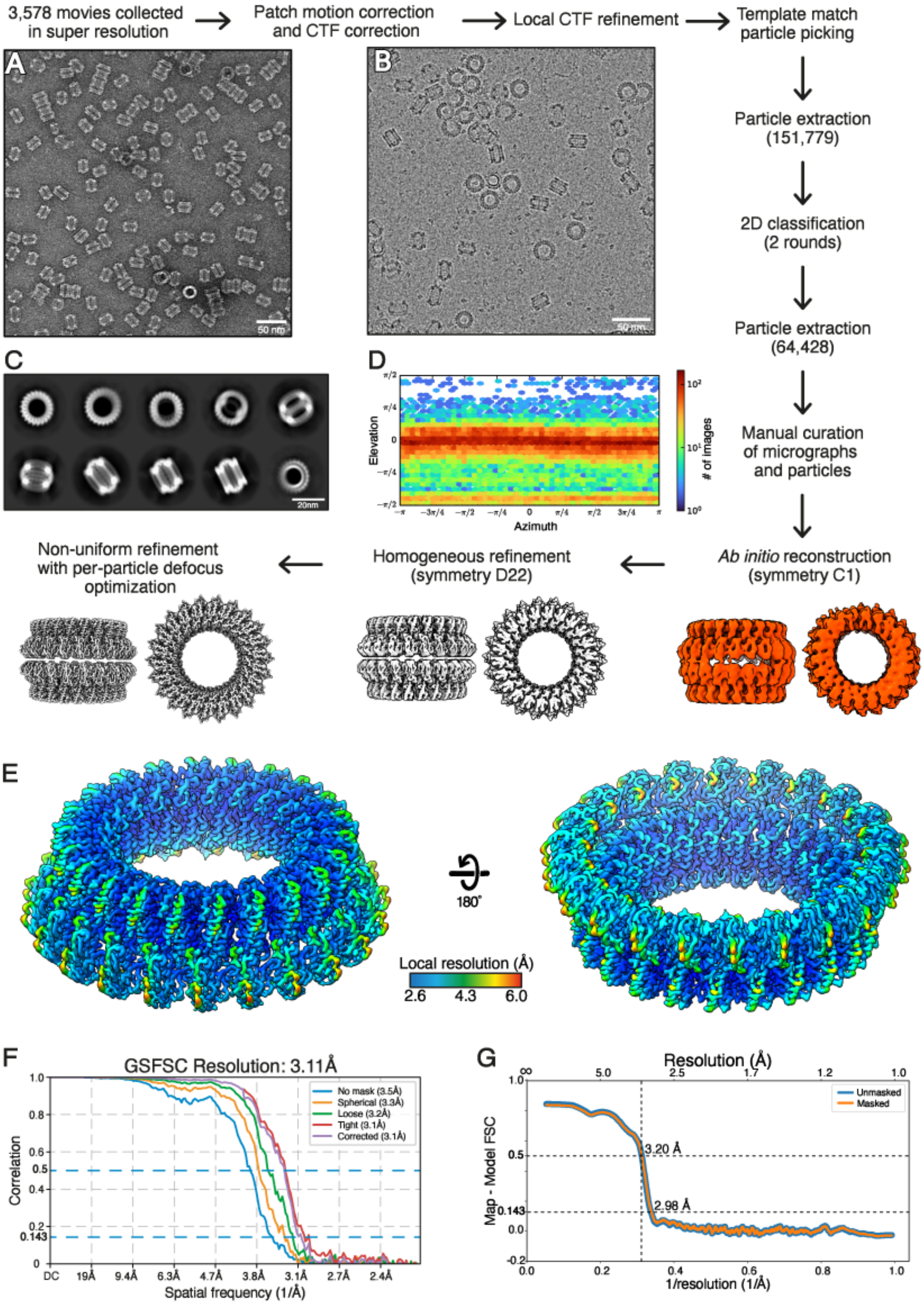
3,578 movies collected in super resolution
Patch motion correction and CTF correction
Local CTF refinement
Template match particle picking
Particle extraction (151,779)
2D classification (2 rounds)
Particle extraction (64,428)
Manual curation of micrographs and particles
Ab initio reconstruction (symmetry C1)
Homogeneous refinement (symmetry D22)
Non-uniform refinement with per-particle defocus optimization
A
B
C
D
E
F
G
50 nm
20nm
Elevation
Azimuth
### of images
Local resolution (Å)
2.6
4.3
6.0
180°
GSFSC Resolution: 3.11Å
No mask (3.5Å)
Spherical (3.3Å)
Loose (3.2Å)
Tight (3.1Å)
Corrected (3.1Å)
Correlation
Spatial frequency (1/Å)
Resolution (Å)
Map - Model FSC
Unmasked
Masked
3.20 Å
2.98 Å
1/resolution (1/Å)

**Fig. S5. Single particle analysis and processing pipeline for the Mpf2Ba1 pre-pore dataset.** (**A, B**) Sample micrographs of pre-pores obtained upon incubation in WCR gut fluid (see Methods) and imaged by negative stain EM (**A**) and cryo-EM (**B**). (**C**) 2D class averages of pre-pores selected for refinement after the last round of classification. (**D**) Plot of the angular distribution of particles contributing to the structure of the pre-pore. (**E**) Local resolution estimation of the pre-pore map. (**F**) FSC curve of the pore showing the final average resolution for the map (3.1 Å at gold standard FSC = 0.143). (**G**) Plot of the map-to-model FSC with and without mask showing very similar FSC curves and resolutions. Details are described in Methods and table S4.

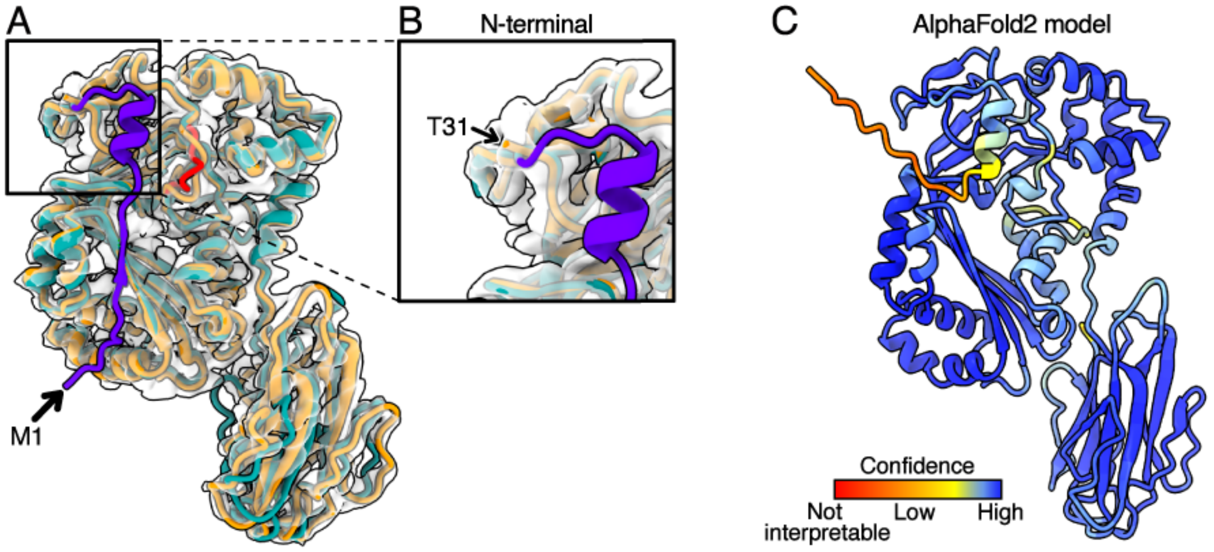


**Fig. S6. Mpf2Ba1 putative N-terminal cleavage site, AlphaFold2 prediction and comparison.** (**A**) Superimposition of Mpf2Ba1 soluble monomer (in cyan) onto pre-pore model (in orange) fitted in the map of a single subunit shows that the first 30 residues of the soluble monomer, from Met1 (black arrow), are absent in the pre-pore map density (in purple). (**B**) Magnified inset showing the density of the pre-pore map starting at N-terminal Thr31 (arrow pointing at Thr31). (**C**) AlphaFold2 (*19*) predicts the N-terminal α-helix but not the short β-strand Val12-Ser15, where those and other residues are predicted with low confidence and not in contact with neighbouring loops/domains. The predicted model has an all-atom RMSD of 2.1 Å and 1.8 Å, for soluble monomer and pre-pore respectively.

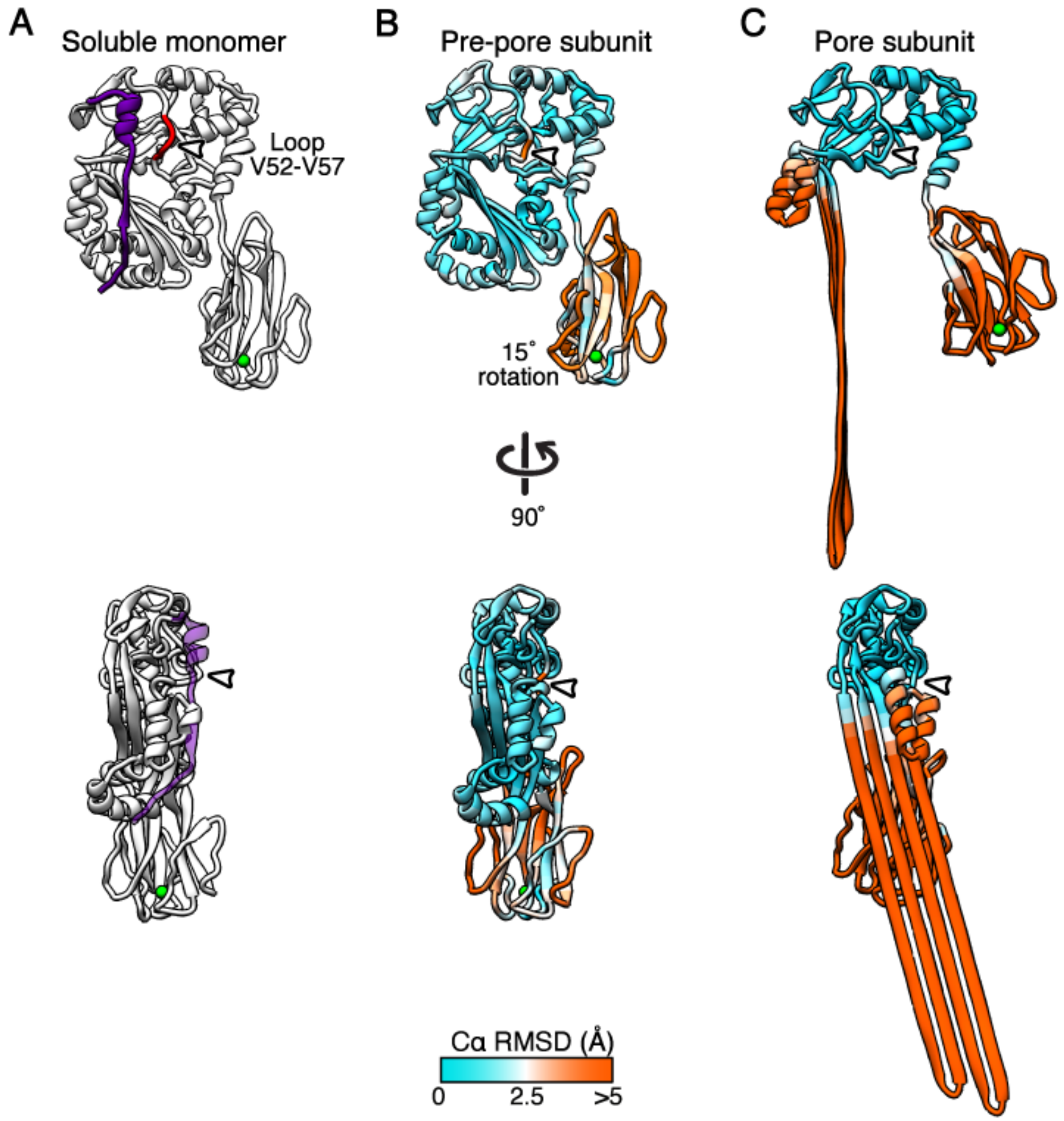


**Fig. S7. Cα RMSD plots of the displacements between Mpf2Ba1 conformations.** (**A**) 30 N-terminal residues in the Mpf2Ba1 soluble monomer (in purple) are not present in the pre-pore and pore models where there is no cryo-EM density to include them; the loop Val52-Val57 is highlighted in red (white arrowhead). (**B**) In the pre-pore the MACPF domain is mostly unchanged (1.2 Å average Cα RMSD for residues 31 to 337, in cyan), except from the loop Val52-Val57 (4.0 Å average Cα RMSD; white arrowheads), and the C-terminal MABP domain that rotates ~15° clockwise relative to the monomeric structure (4.2 Å average Cα RMSD for residues 338 to 484). (**C**) In the pore structure, the upper part of the MACPF domain remains unchanged (1.4 Å Cα RMSD) while TMHs undergo a big conformational change into β-hairpins, and C-terminal domain and HTH motif move ~12 Å and ~8 Å respectively from their center of mass compared to the pre-pore conformation. The Cα RMSD distances were calculated between soluble monomer and pre-pore, reported in (**B**), and between pre-pore and pore, reported in (**C**).

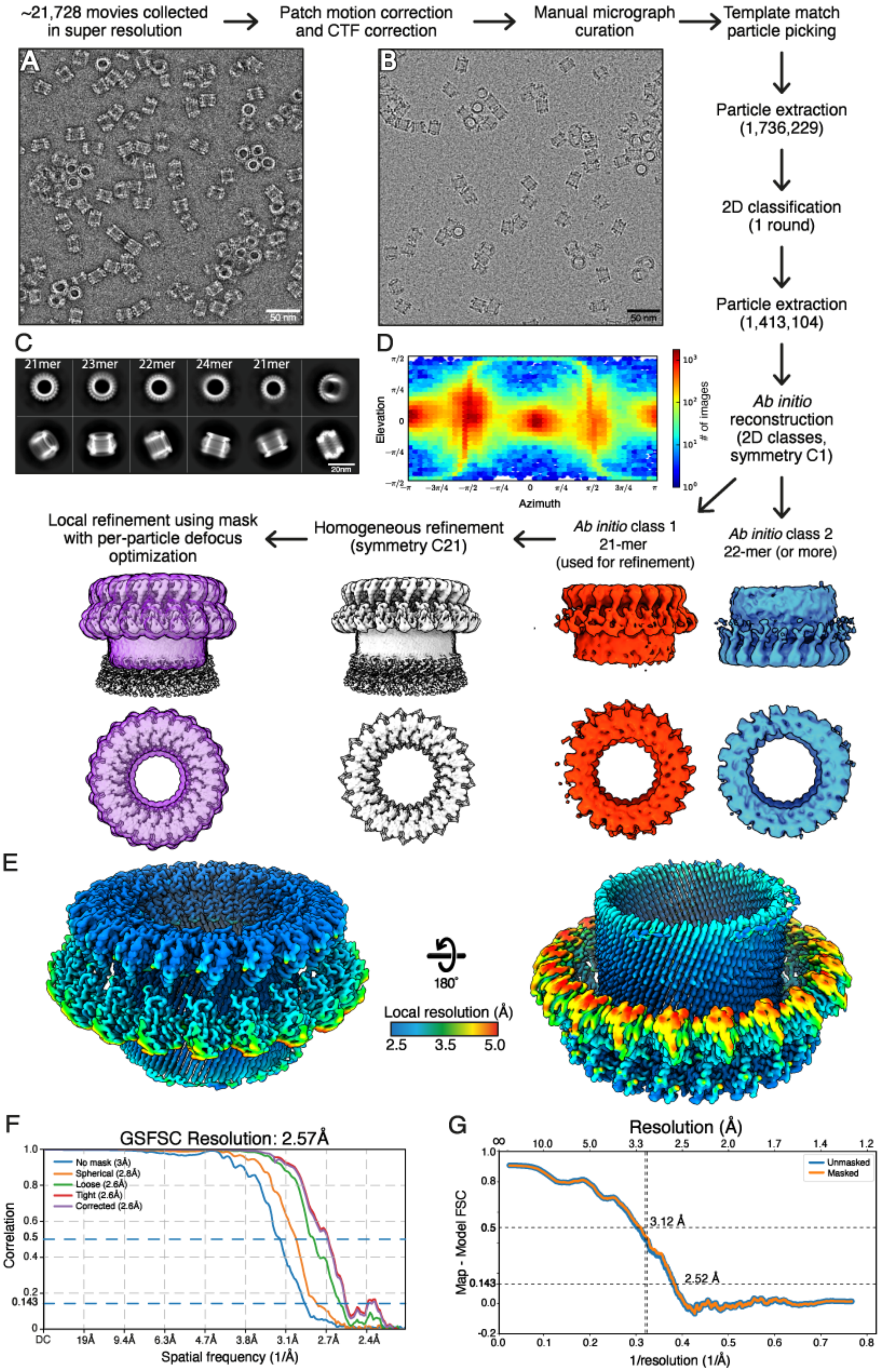
~21,728 movies collected in super resolution
Patch motion correction and CTF correction
Manual micrograph curation
Template match particle picking
Particle extraction (1,736,229)
2D classification (1 round)
Particle extraction (1,413,104)
Ab initio reconstruction (2D classes, symmetry C1)
Ab initio class 1 21-mer (used for refinement)
Ab initio class 2 22-mer (or more)
Homogeneous refinement (symmetry C21)
Local refinement using mask with per-particle defocus optimization
A
B
50 nm
C
21mer
23mer
22mer
24mer
21mer
20nm
D
Elevation
Azimuth
### of images
E
180°
Local resolution (Å)
2.5
3.5
5.0
F
GSFSC Resolution: 2.57Å
No mask (3Å)
Spherical (2.8Å)
Loose (2.6Å)
Tight (2.6Å)
Corrected (2.6Å)
Correlation
Spatial frequency (1/Å)
G
Resolution (Å)
Map - Model FSC
1/resolution (1/Å)
Unmasked
Masked
3.12 Å
2.52 Å

**Fig. S8. Single particle analysis and processing pipeline for the Mpf2Ba1 pore dataset.** (**A, B**) Sample micrographs of pores obtained upon incubation at ~53˚C (see Methods), imaged by negative stain EM (**A**) and cryo-EM (**B**). (**C**) 2D class averages of pores obtained at the last round of classification showed a distribution of oligomer stoichiometries, ranging from the most populated 22-fold symmetry (52% of end view particles) to 21-fold (30%), 23-fold (11%) and 24-fold (7%), demonstrating heterogeneity of oligomerization. The mask created for the local refinement to enhance the density of the 21-mer pore is shown in violet. (**D**) Plot of the angular distribution of particles contributing to the structure of the pore. (**E**) Local resolution estimation of the pore map. (**F**) FSC curve of the pore showing the final average resolution for the map (2.6 Å at FSC = 0.143). (**G**) Plot of the map-to-model FSC with and without mask showing very similar FSC curves and resolutions. Details are described in Methods and table S4.

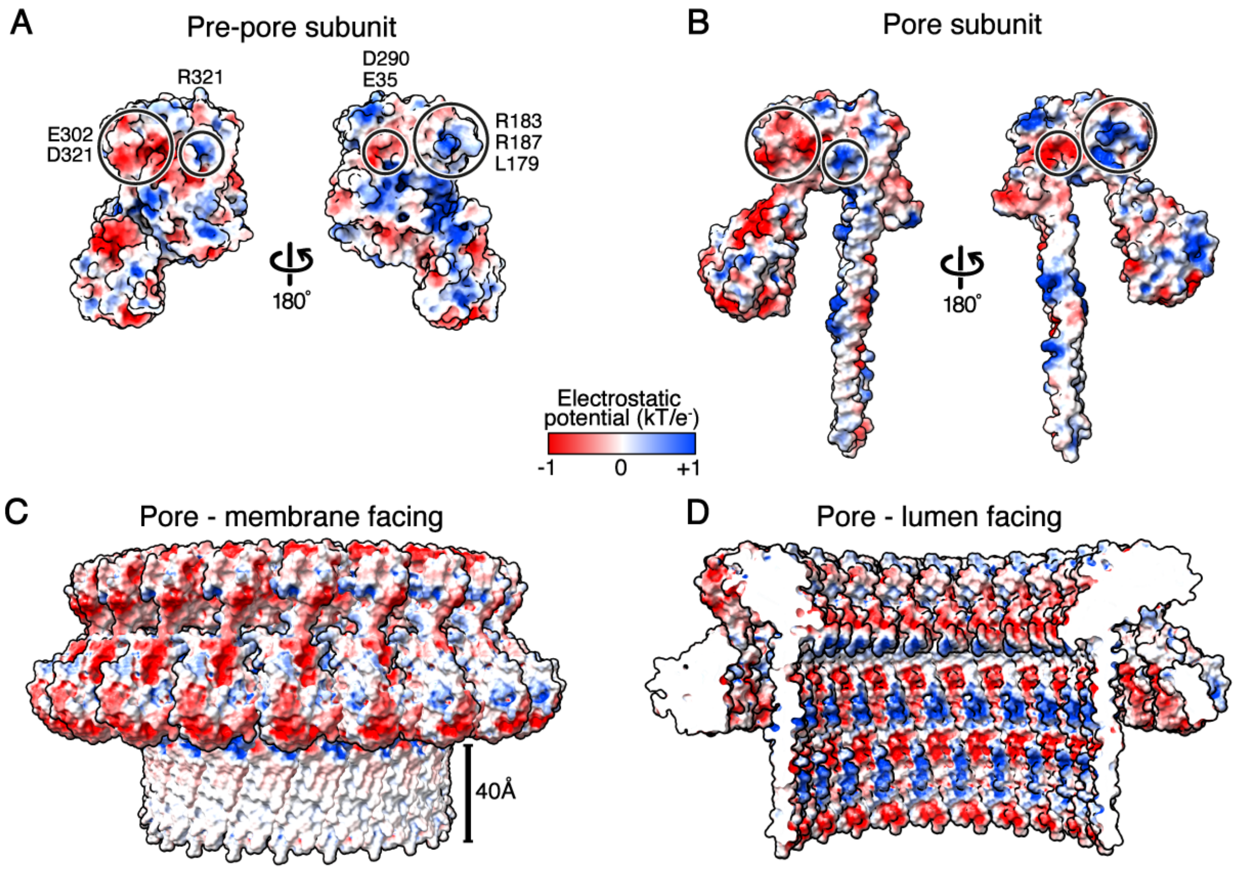


**Fig. S9. Analysis of the electrostatic potential maps of Mpf2Ba1 pre-pore and pore**. (**A, B**) MACPF domains of neighbouring pre-pore and pore subunits show opposite charges in complementary sites of interaction: on the left, the negative pocket of Glu302, Asp321 and the positive charge of Arg151 match exactly with the positive pocket of Lys179, Arg183 and Arg187, and the negative pocket of Glu35 and Asp290, on the right (black circles). (**C**) The distribution of charges for the entire pore structure shows the membrane-facing region with a charge-neutral belt, ~40 Å in height, enriched in Phe, Tyr, Val and Ile hydrophobic residues, while (**D**) the lumen-facing part is highly polar and includes charged residues such as Glu, Asp, Lys and up to 9 Serines per subunit.

| | Amino acid position | | | | | | | | |
|---|---|---|---|---|---|---|---|---|---|
| | **81** | **92** | **100** | **322** | **334** | **340** | **347** | **435** | **454** |
| **Mpf2Ba1 WT** | I | Y | I | Y | Y | Y | I | Y | I |
| **Mpf2Ba1-1167** | L | F | L | F | F | F | L | F | L |

**Table S1.** Conservative mutations of Mpf2Ba1-1167 relative to the wild-type protein.

| | $LC/IC_{50}$ | WCR (ppm) | NCR (ppm) | ***Diabrotica speciosa*** **(ppm)** |
|---|---|---|---|---|
| **Mpf2Ba1 WT** | $LC_{50}$ | 16.3 | 35.6 | >400* |
| | $IC_{50}$ | 7.4 | 13 | 320 |
| **Mpf2Ba1-1167** | $LC_{50}$ | 6.49 | ND | ND |
| | $IC_{50}$ | 4.88 | ND | ND |

**Table S2.** Mpf2Ba1 wild type and Mpf2Ba1-1167 bioactivity in artificial diet bioassays ($IC_{50}$ and $LC_{50}$) against *Diabrotica virgifera* (WCR), *Diabrotica barberi* (NCR) and *Diabrotica speciosa.* *Mortality was 35% at 400 ppm dose.

| Protein | WCR Population | $LC_{50}$, µg/ml | 95% Confidence Interval | Slope ± Standard Error | n | Resistance Ratio (RR) |
|---|---|---|---|---|---|---|
| **Gpp34Ab1/Tpp35Ab1** | Susceptible | 5.90 | 3.06 - 8.60 | 3.27 ± 0.910 | 144 | 8 |
| | Readlyn | 45.7 | 23.6 - 85.3 | 1.08 ± 0.207 | 144 | |
| **mCry3A** | Susceptible | 6.97 | 1.61 - 13.7 | 1.90 ± 0.600 | 140 | 43 |
| | Readlyn | 302 | 87.1 - 6680 | 0.644 ± 0.230 | 212 | |
| **Mpf2Ba1 WT** | Susceptible | 10.6 | 8.22 - 13.6 | 2.34 ± 0.277 | 287 | 0.7 |
| | Readlyn | 7.46 | 5.62 - 9.72 | 2.32 ± 0.290 | 288 | |
| **Mpf2Ba1-1167** | Susceptible | 14.3 | ND | ND | 144 | 0.8 |
| | Readlyn | 11.4 | 6.80 - 16.9 | 2.07 ± 0.397 | 144 | |

**Table S3.** Bioactivity of Mpf2Ba1 and Mpf2Ba1-1167 against susceptible and Readlyn populations of WCR. Artificial diet bioassay data indicate that Mpf2Ba1 and Mpf2Ba1-1167 are not cross-resistant to the Cry3A and Gpp34Ab1/Tpp35Ab1 proteins. Resistance ratio (RR) defined as the $LC_{50}$ value determined for resistant WCR divided by the $LC_{50}$ value determined for susceptible WCR. ND: could not determine.

| Cryo-EM imaging | Pre-pore | Pore |
|---|---|---|
| Magnification | ×130k | ×81k |
| Voltage [kV] | 300 | 300 |
| Direct Electron Detector | Gatan K2 | Gatan K3 |
| TEM mode | EFTEM nanoprobe | EFTEM nanoprobe |
| Energy filter slit width | 20 eV | 20 eV |
| Exposure rate [$e^-$/pix/s] | 5.70 | 19.22 |
| Exposure rate [$e^-$/Å$^2$/s] | 4.95 | 16.88 |
| Exposure time [s] | 8 | 3 |
| Total exposure [$e^-$/Å$^2$] | 39.56 | 50.65 |
| Movie frames | 40 | 50 |
| Exposure per frame [e/Å$^2$/frame] | 0.99 | 1.01 |
| Target defocus [µm] | 1.5-3.0 | 1.5-3.3 |
| Pixel size [Å] | 1.05 | 1.067 |
| Super resolution | Yes | Yes |
| Total movies | 3,578 | 21,728 |
| **Single Particle Analysis** | | |
| Initial particle images | 151,779 | 1,736,229 |
| Final particle images | 64,248 | 1,413,104 |
| Symmetry imposed | D22 | C21 |
| Map resolution [Å] | 3.3 Å | 3.0 Å |
| FSC threshold | 0.143 | 0.143 |

**Table S4.** Cryo-EM imaging and single particle analysis parameters used for pre-pore and pore dataset collection and 3D reconstructions.

A

| Validation scores | Pre-pore | Pore |
|---|---|---|
| Q-score (σ = 0.6) | 0.64 | 0.59 |
| Average SMOCf score | 0.85 | 0.88 |
| Strudel score (2.8-3.0 Å motif library) | 0.946 | 0.895 |
| MolProbity score | 1.67 | 1.54 |

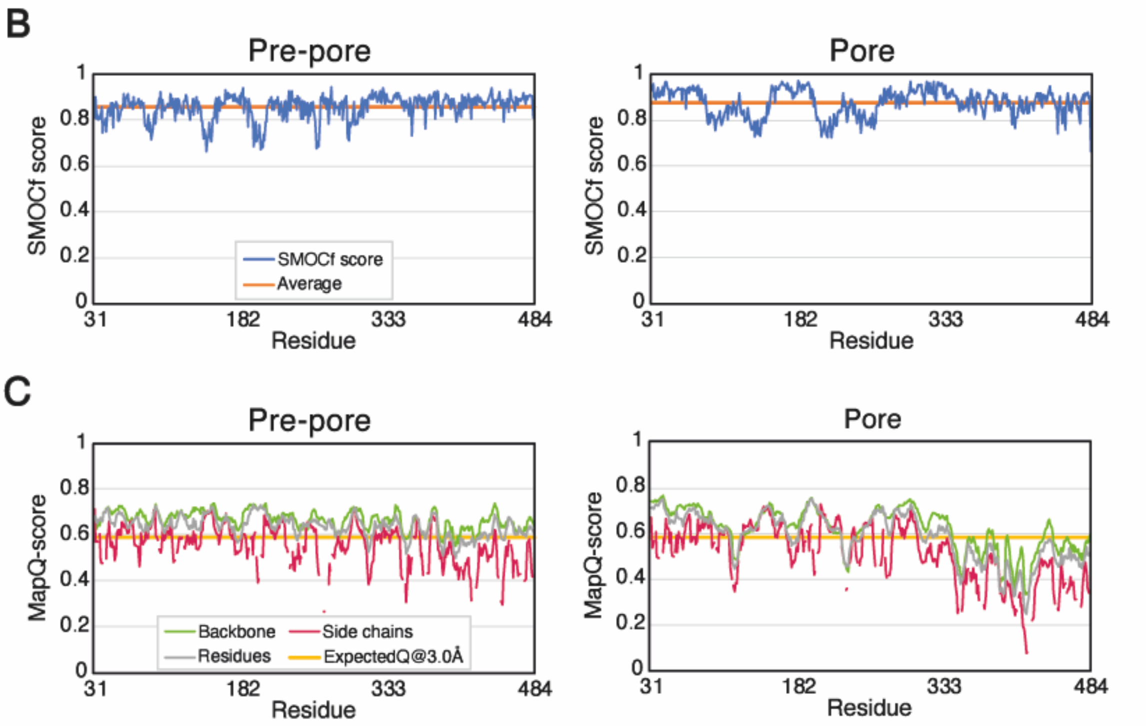


**Table S5. Table and plots of pre-pore and pore model validation scores.** (**A**) Final validation scores for the pre-pore and pore models calculated using different refinement and validation programs. (**B**) SMOCf scores (*62, 71*) are shown per residue in the plots. (**C**) Backbone, side chains and residues Q-scores from MapQ (70) are calculated with the default width value of σ = 0.6 and showed averaged over 3 residues in the plots.

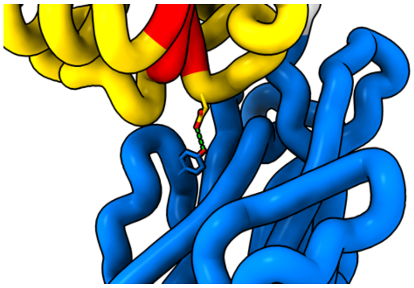

**Movie S1.**
Glu137-Tyr460 H-bond breakage.

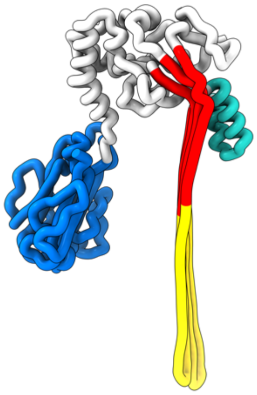

**Movie S2.**
Conformational changes of a single Mpf2Ba1 subunit from crystal to pre-pore to pore.

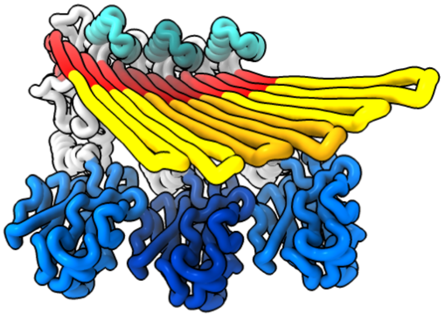

**Movie S3.**
Conformational change from pre-pore to pore of three neighbouring Mpf2Ba1 subunits to show the rearrangement of TMHs and HTH.
